# Redistribution of cholesterol from vesicle to plasmalemma controls fusion pore geometry

**DOI:** 10.1101/2020.04.06.027169

**Authors:** Boštjan Rituper, Alenka Guček, Marjeta Lisjak, Urszula Gorska, Aleksandra Šakanović, Saša Trkov Bobnar, Eva Lasič, Mićo Božić, Prabhodh S. Abbineni, Jernej Jorgačevski, Marko Kreft, Alexei Verkhratsky, Frances M. Platt, Gregor Anderluh, Matjaž Stenovec, Bojan Božič, Jens R. Coorssen, Robert Zorec

**Affiliations:** Laboratory of Neuroendocrinology-Molecular Cell Physiology, Institute of Pathophysiology, University of Ljubljana, Faculty of Medicine, Ljubljana, Slovenia; Celica Biomedical, 1000, Ljubljana, Slovenia; Laboratory for Molecular Biology and Nanobiotechnology, National Institute of Chemistry, Ljubljana, Slovenia; Department of Pharmacology, University of Michigan, Ann Arbor, MI 48109-5632, USA; Faculty of Biology, Medicine and Health, The University of Manchester, Manchester, M13 9PT, UK; Achucarro Center for Neuroscience, IKERBASQUE, 48011 Bilbao, Spain; Department of Pharmacology, University of Oxford, Oxford, OX1 3QT, UK; Institute of Biophysics, Faculty of Medicine, University of Ljubljana, Slovenia; Department of Health Sciences, Faculty of Applied Health Sciences and Department of Biological Sciences, Faculty of Mathematics & Science, Brock University, St Catherine’s, Ontario, Canada; Department of Medical Cell Biology, Uppsala University, Uppsala, Sweden; Smoluchowski Institute of Physics, Jagiellonian University, Krakow, Poland

## Abstract

Eukaryotic vesicles fuse with the plasmalemma to form the fusion pore, previously considered to be unstable with widening of the pore diameter. Recent studies established that the pore diameter is stable, reflecting balanced forces of widening and closure. Proteins are considered key regulators of the fusion pore, whereas the role of membrane lipids remains unclear. Super-resolution microscopy revealed that lactotroph secretory vesicles discharge cholesterol after stimulation of exocytosis; subsequently, vesicle cholesterol redistributes to the outer leaflet of the plasmalemma. Cholesterol depletion in lactotrophs and astrocytes evokes release of vesicle hormone, indicating that cholesterol constricts the fusion pore. A new model of cholesterol-dependent fusion pore diameter regulation is proposed. High-resolution measurements of fusion pore conductance confirmed that the fusion pore widens with cholesterol depletion and constricts with cholesterol enrichment. In fibroblasts lacking the Npc1 protein, in which cholesterol accumulates in vesicles, the fusion pore is narrower than in controls, showing that cholesterol regulates fusion pore geometry.

**Graphical Abstract:** 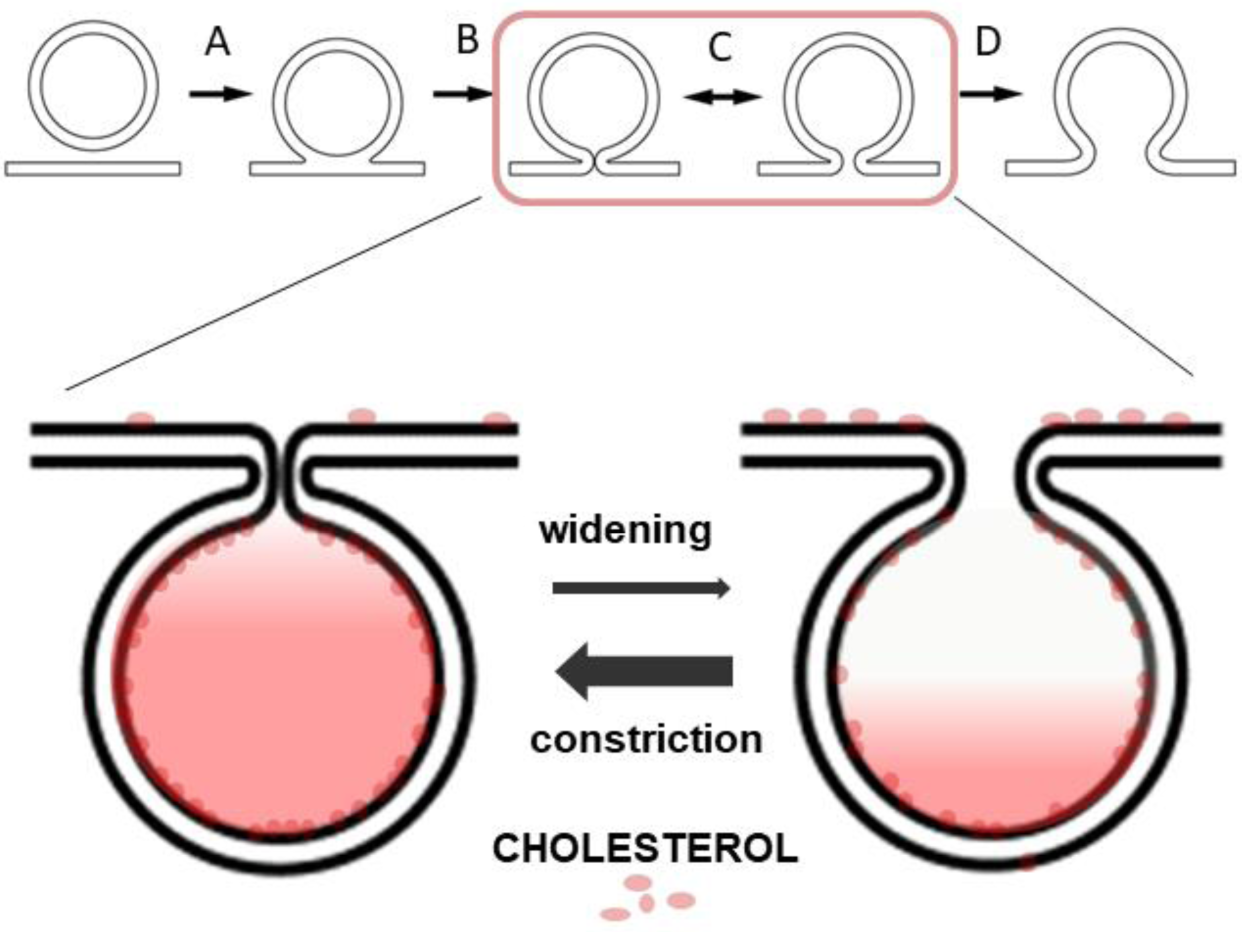

Top: stages through which a vesicle interacts with the plasmalemma. Stage A denotes hemifusion, which proceeds to stage B, with a narrow fusion pore, which can then reversibly open (stage C), before widening fully (stage D). Bottom: redistribution of cholesterol from the vesicle to the outer leaflet of the plasmalemma controls fusion pore constriction.

**In Brief:** A membrane pore is formed when the vesicle membrane fuses with the plasmalemma. Proteins were considered key regulators of the opening and closing of this fusion pore. Here, evidence is provided to show that cholesterol, a membrane constituent, determines a radial force constricting the fusion pore, revealing that the fusion pore functions as a proteolipidic structure.

**Highlights:**

- Intravesicular cholesterol redistributes to the outer leaflet of the plasmalemma.
- Cholesterol depletion widens the fusion pore, whereas cholesterol enrichment constricts the fusion pore.
- A model of cholesterol-dependent force preventing fusion pore widening is developed.
- Disease-related increase in vesicle cholesterol constricts the fusion pore.

## INTRODUCTION

Membrane fusion, fundamental to the life of eukaryotic cells, contributes to diverse processes, including catabolic and biosynthetic pathways, as well as to vesicle-based secretion (Brose et al., 2019). Regulated exocytosis, which proceeds through a merger between the vesicle membrane and the plasmalemma (PM), leads to the formation of a fusion pore, through which molecules stored in the vesicle lumen, including hormones and neurotransmitters, exit into the extracellular space (Sharma and Lindau, 2018). The views that emerged decades ago (Rand and Parsegian, 1986), assumed that once the fusion pore forms, the vesicle membrane integrates abruptly and fully with the PM (full exocytosis). However, recent studies have postulated that the open fusion pore may close again after a while (transient or reversible exocytosis). The stages that a vesicle undergoes during exocytosis were confirmed by measurements of discrete steps in membrane capacitance (C_m_), which is linearly related to the membrane area, and is affected by vesicle fusion and fission (Neher and Marty, 1982). Advancements through the stages of vesicle-PM fusion appear to depend on vesicle size (Flasker et al., 2013; Jorgacevski et al., 2010; Shin et al., 2018). In synaptic communication, that is specialized for fast signaling, vesicles with relatively small diameters predominantly undergo full exocytosis (Grabner and Moser, 2018; He et al., 2006; Klyachko and Jackson, 2002). The fusion pore opening follows the calcium trigger by a millisecond and is considered to be driven by the soluble N-ethylmaleimide-sensitive factor attachment protein receptors (SNAREs), a protein complex responsible for facilitating exocytotic membrane fusion (Brose et al., 2019).

The structure of the fusion pore walls is still debated, although it is widely considered to be proteolipidic in nature (Sharma and Lindau, 2018).Therefore, it is likely that both proteins and lipid membrane molecules affect the fusion pore, including stabilizing its structure. In this study, we examined whether cholesterol, the most common steroid in humans and a major constituent of the cell membrane, affects fusion pore geometry. To this end, we monitored single-vesicle cargo discharge and characterized unitary exocytotic events by measuring changes in C_m_ (Akerman et al., 1991; Mason et al., 1988; Neher and Marty, 1982; Sikdar et al., 1989; Zorec et al., 1991). Previous studies indicated that cholesterol may regulate exocytosis, most likely by affecting the local curvature of the membrane to promote merger of the apposed monolayers (Churchward and Coorssen, 2009; Churchward et al., 2005a; Cookson et al., 2013; Rituper et al., 2013a; Rituper et al., 2012; Wang et al., 2010). Another study demonstrated that cholesterol facilitates spontaneous and inhibits evoked synaptic secretion (Wasser et al., 2007). In endocrine pituitary cells, cholesterol appears to be necessary for the fusion of vesicle and plasma membranes (Rituper et al., 2013a; Rituper et al., 2012), whereas in insulin-secreting cells, excess cholesterol inhibits vesicle exocytosis (Xu et al., 2017). These contradictory results may arise from cholesterol acting differently at various stages of vesicle interactions with the PM.

Using a cholesterol marker (Lasic et al., 2019; Skocaj et al., 2014), we found that during stimulated exocytosis, vesicle cholesterol mixes with the PM of the outer leaflet; this occurs by cholesterol transfer through the fusion pore, likely driven by the cholesterol concentration gradient between these two membrane compartments. Here, depleting cholesterol by exposing cells to methyl-β-cyclodextrin (MβCD) stimulated vesicle cargo release independently of cytosolic Ca^2+^, indicating that cholesterol may contribute to the constriction of the fusion pore. We propose a model of cholesterol-dependent radial force that regulates the fusion pore diameter. High-resolution measurements of C_m_ and fusion pore conductance (G_p_), which reports the electrical geometry of the fusion pore, confirmed our theoretical predictions that cholesterol depletion widens and cholesterol enrichment constricts the fusion pore. Consistent with this hypothesis, cholesterol-rich vesicles in fibroblasts lacking the Niemann-Pick Type C1 (Npc1) protein, a cellular model of the lysosomal storage disease Niemann-Pick disease Type C1 (NPC), exhibited a narrower fusion pore than wild-type controls. Our findings provide a conceptual advancement of our understanding of membrane fusion pores.

## RESULTS

### Secretory Vesicles Contain Cholesterol

Lactotroph vesicles store and secrete prolactin (PRL). We labeled these cells by anti-PRL antibodies (Figure 1A) together with fluorescent (TopFluor) cholesterol (Figure 1B), which has similar properties to native cholesterol (Holtta-Vuori et al., 2008), to determine whether PRL and cholesterol are stored in the same vesicles. The exposure of cells to TopFluor for minutes predominantly labeled the PM (Rituper et al., 2013a). Longer exposure, that ended with a 2 min wash of cells with MβCD to remove background fluorescence (see STAR Methods), led to most vesicles containing both markers (Figure 1C). Diameters of structures with both markers (Figures 1D–1F) were determined by constructing intensity profiles of labeled structures, fitting Gaussian curves and measuring full width at half maximum (FWHM = 2σ(2ln2)^1/2^, where σ denotes standard deviation) (Figure 1C, inset). The PRL- and TopFluor-positive structures (n = 384, from ten cells) were of similar diameter: 249 ± 2 nm (Figure 1D) and 243 ± 3 nm (Figure 1E), respectively. Fitting the pairs of PRL- and TopFluor-positive vesicle diameters by linear regression yielded a slope of k = 0.98 ± 0.01 (correlation coefficient r = 0.84, *p* < 0.001, n = 384; Figure 1F), indicating that cholesterol may be present in PRL-containing secretory vesicles, consistent with previous observations (Bogan et al., 2012). This suggests that stimulation of regulated exocytosis may instigate the transfer of vesicle cholesterol, thus affecting the composition of cholesterol in the PM.

**Figure 1.**
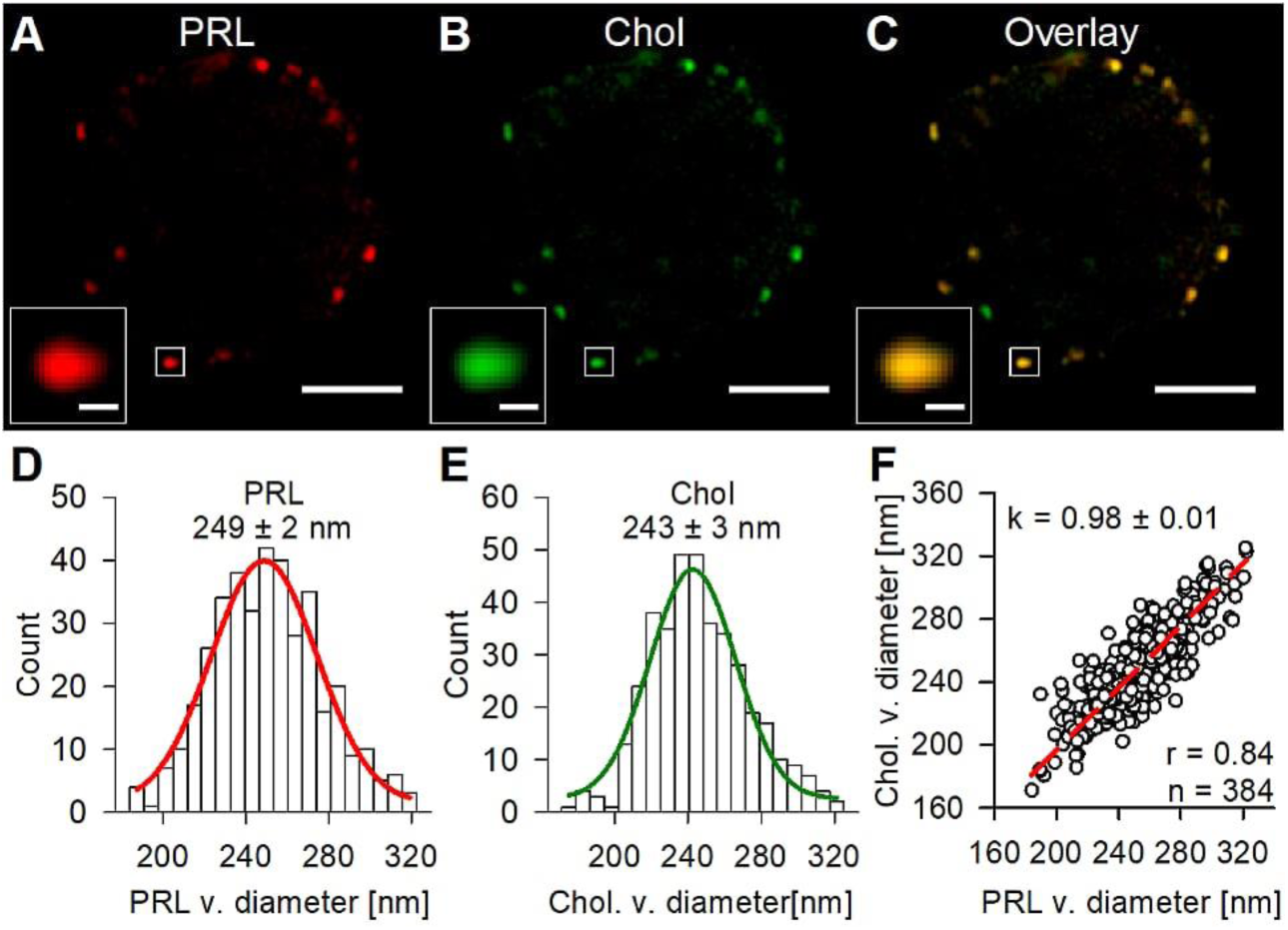
Secretory Vesicles Contain Prolactin (PRL) and Cholesterol (Chol) (A–C) Deconvoluted images of a lactotroph, stained by an anti-PRL antibody (red, A) and fluorescent TopFluor-labeled cholesterol (green, B). (C) PRL and Chol signals are overlaid (colocalized pixels in yellow). Insets in the lower left corners (A–C) represent colocalized PRL and Chol in a vesicle. (D) Frequency distribution (count) of PRL vesicles versus their diameter (PRL v. diameter). The red line represents the fitted Gaussian (see also Figure S1) with a mean PRL vesicle diameter of 249 ± 2 nm (n = 384; bin width, 10 nm; ten cells). (E) Frequency distribution (count) of Chol vesicles versus their diameter. The green line depicts the fitted Gaussian with a mean vesicle diameter of 243 ± 3 nm (n = 384; bin width, 10 nm; ten cells). (F) PRL vesicle (v.) diameter plotted versus Chol vesicle (v.) diameter. The red line depicts a linear regression (passing through the origin) of the form: Chol. v. diameter = k × PRL v. diameter; correlation coefficient, r = 0.84 (p < 0.001); the slope of the line k = 0.98 ± 0.01 (in red; p < 0.001). Scale bar, 3 µm; inset scale bar, 200 nm in (A)–(C).

### Stimulation of Regulated Exocytosis Reduces the Density and Size of Cholesterol-Rich PM Domains

To test how stimulated exocytosis affects cholesterol-rich PM domains, we first used a cholesterol-specific marker, mCherry-D4-PFO (D4) (Lasic et al., 2019; Maekawa, 2017; Ohno-Iwashita et al., 1990; Skocaj et al., 2014), to visualize the cholesterol content in the PM, which is rich in cholesterol in pituitary cells (Rituper et al., 2013a). Cells were exposed to D4, washed subsequently with extracellular solution (vehicle, protocol A, Table S1), and finally fixed with 2% formaldehyde. The D4 labelled the PM (Figure 2A). Neither intracellular injection of the D4 peptide nor transfection of cells with the mCherry-D4-encoding plasmid produced significant labeling of the cytoplasmic leaflet of the PM (Figure S1). In MβCD-treated (10 mM, 30 min) cells (Najafinobar et al., 2016; Rituper et al., 2013a), the PM D4 staining was strongly reduced (Figure 2B, right), indicating, together with the results in Figure S1, that D4 associates with the cholesterol-rich domains in the outer PM leaflet. The extent of D4 labeling was determined by delineating a band overlaying the PM (Figure S2) and calculating the percentage of D4-positive pixels, relative to all the pixels in the region of interest. MβCD reduced the extent of D4 labeling from 9.0% ± 0.5% in controls to 2.1% ± 0.3% in MβCD-treated cells (*p* < 0.001, n = 50 optical slices from five cells per group). This is consistent with the biochemically determined cholesterol content (Amplex Red, see STAR Methods) of MβCD-treated pituitary cells, which was 56.8% ± 10.6%, normalized to controls; a similar reduction to 52.3% ± 4.9% was also obtained in astrocytes (Table S2). MβCD selectively extracted cholesterol from cells without affecting other membrane lipids, as quantified by high-performance thin layer chromatography (Table S2).

**Figure 2.**
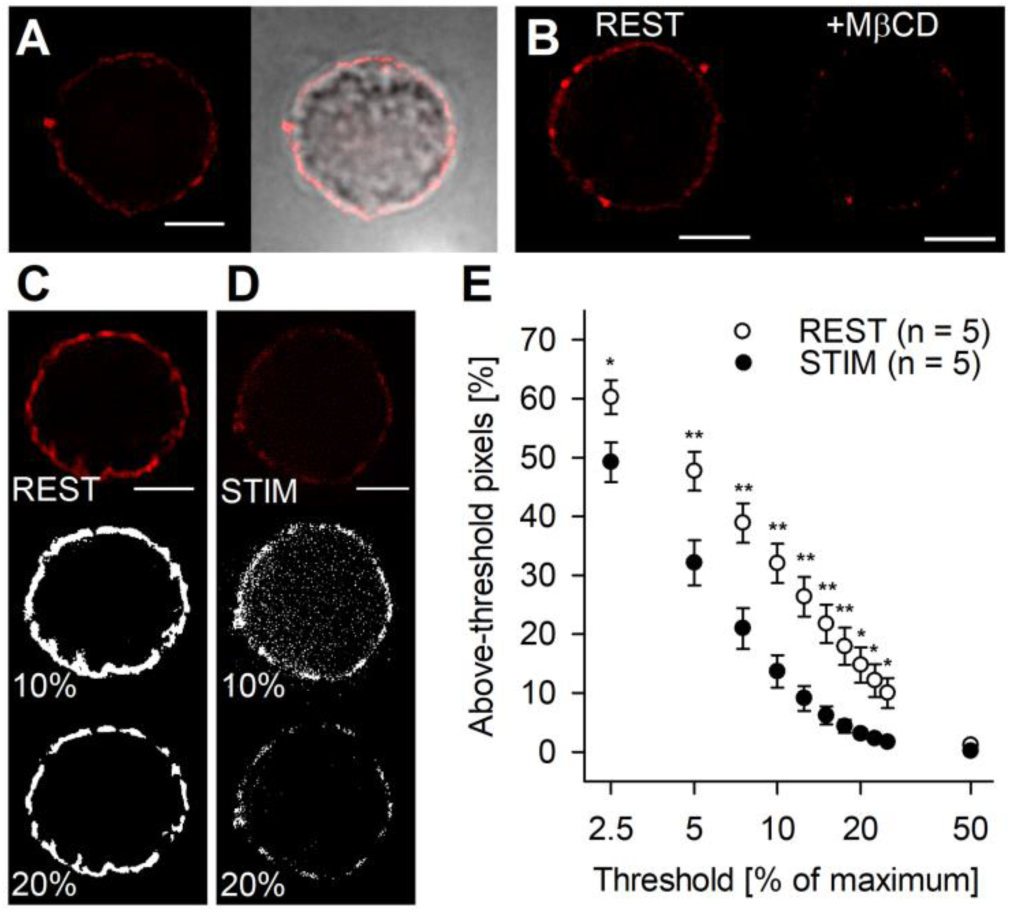
Stimulation of Regulated Exocytosis Decreases the Density of Cholesterol-Rich Domains in the Outer Leaflet of the Plasmalemma, Labeled by mCherry-D4-PFO (D4) in Lactotrophs. (A) Super-resolution (left) and transmission (right) images of a D4-labeled lactotroph (protocol A; Table S2), revealing the density of cholesterol-rich membrane domains in the outer leaflet of the PM. (B) D4-labeled lactotroph at rest (REST, left) and after cholesterol extraction with methyl-β-cyclodextrin (+MβCD, 10 mM, 30 min, right). (C and D) Representative structured illumination microscopy images of (C) D4-labeled resting (REST, labeling protocol A) and (D) stimulated lactotrophs (STIM; 50 mM KCl, labeling protocol B) and thresholded mask images (at 10% and 20% threshold, in white. See STAR Methods, Figure S1, and Table S2. (E) The above-threshold membrane D4-positive pixels (expressed as a percentage of all pixels in the region of interest) versus the threshold (percentage of maximum) in resting (white circles) and stimulated (black circles) cells. Asterisks denote a significant reduction in D4-positive above-threshold pixels in the plasma membrane of stimulated (STIM) versus resting (REST) lactotrophs (n > 30 optical slices from five cells per group). Data are presented as means ± SEM in (E). **p < 0.01, *p < 0.05. Student’s t-test in (E). Scale bars, 3 μm in (A)–(D).

Second, we tested whether the distribution of cholesterol in the PM was affected by regulated exocytosis. D4-labeled cells were depolarized with extracellular solution containing 50 mM KCl, which triggers exocytotic secretion (Stenovec et al., 2004); and were subsequently examined by microscopy (Table S1, protocol B). D4-labeling of the PM was reduced in the stimulated cells (Figures 2C and 2D). As the decrease in the signal depends on the noise level, we assessed the changes in D4 area at several intensity thresholds (middle and bottom panels, Figures 2C and 2D). The D4-labeled pixels above the threshold were similarly significantly reduced in exocytosis-stimulated cells (Figure 2E; n = 30 optical slices, five cells), indicating that the D4-unlabeled cholesterol delivered from vesicles diluted the D4-labeled cholesterol in the PM. By readjusting the acquisition parameters for faint signals, we further tested the density reduction of D4-positive domains after stimulation of exocytosis (p < 0.001, Figures S3A and S3B), and also determined that the size of individual D4 domains (insets in Figures 2C and 2D) decreased after stimulation of regulated exocytosis (*p* < 0.05, Figures S3C and S3D).

These data indicate that stimulation of regulated exocytosis leads to changes in cholesterol distribution in the external leaflet of the PM.

### Vesicle Cholesterol Delivery to the PM Correlates with Exocytotic Release of Prolactin

To further examine whether regulated exocytosis initiates cholesterol redistribution to the PM, we performed three sets of experiments using distinct labeling approaches (protocols C, D, and E; Table S1).

First, cholesterol-rich domains were labeled with D4 and PRL at the PM surface with antibodies (Figures 3A and 3B). In controls (REST), protocol C (Table S1) was used to label vesicle-stored PRL and cholesterol-rich domains in non-stimulated cells; protocol D (Table S1) was used to label surface-deposited PRL granules in stimulated cells, and cholesterol-rich domains were labeled before stimulation of exocytosis.

**Figure 3.**
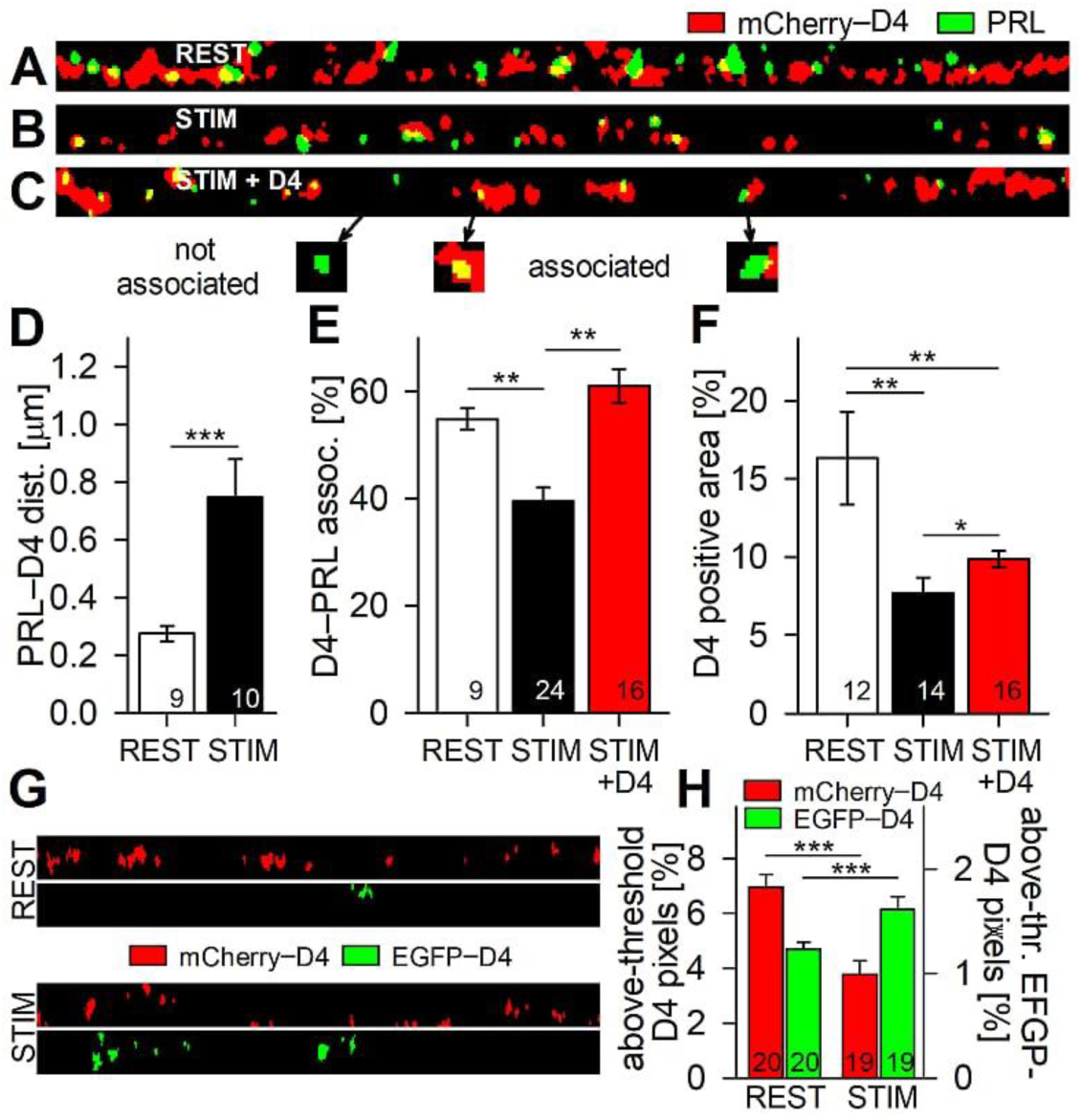
Vesicle Cholesterol Delivery to the Plasmalemma (PM) is Concomitant with Vesicule Prolactin (PRL) Release during Stimulated Exocytosis. (A) Straightened thresholded structured illumination microscopy image of the PM cell region in a resting lactotroph (REST, labeling protocol C). Red, mCherry-D4 (D4); green, PRL vesicles; yellow, colocalized D4- and PRL-positive pixels. See STAR Methods, Figure S1, and Table S2. Note the colocalization between D4 and PRL signals, indicating association of cholesterol-rich domains (D4) with released PRL vesicle cargo. (B) Straightened thresholded structured illumination microscopy image of the PM cell region in a stimulated lactotroph (STIM; 50 mM KCl, labeling protocol D). Red, D4; green, PRL surface-attached PRL granules. Note the reduced colocalization between D4 and PRL signals, indicating reduced association of cholesterol-rich domains (D4) with PRL cargo deposited on the cell surface after regulated exocytosis. (C) Straightened thresholded structured illumination microscopy image of the PM cell region in a stimulated lactotroph (STIM + D4; with 50 mM KCl, labeling protocol E), in which D4-labeling solution was also present during PRL labeling (see Table S2). Insets present examples of associated D4 and PRL structures, defined as either overlaid or in contact. Note that this experiment is similar to that in (B), exept that when stimulation was administered, the D4 marker was present, indicating an increased association of cholesterol-rich domains (D4) with PRL cargo deposited on the surface of cells after regulated exocytosis. (D) The shortest distance between the D4 domain and the nearest PRL-positive structure (PRL–D4 dist., in µm) in resting (REST) and stimulated (STIM) cells. (E) D4 to PRL association (D4–PRL assoc., presented as the percentage of associated D4 structures versus all D4 structures). Note that stimulation of regulated exocytosis (STIM) reduces the association of D4 domains with vesicle cargo PRL, likely due to the incorporation of vesicle cholesterol into the PM. (F) Relative D4-labeled membrane area (percentage of above-threshold pixels relative to all pixels in the PM region) in resting (REST) and stimulated (STIM and STIM + D4) cells. (G) Upper panel: Representative straightened thresholded image of the PM region of a resting lactotroph (REST), labeled with D4 (upper band) followed by labeling with EGFP-D4-PFO (EGFP-D4, lower band). Lower panel: Representative straightened thresholded image of the PM region of a D4-labeled lactotroph (upper band) followed by labeling with EGFP-D4 after stimulation (STIM) of regulated exocytosis with 50 mM KCl. Note that stimulation of regulated exocytosis increased the EGFP-D4 labeling, whereas D4 labeling was decreased, consistent with the formation of additional cholesterol-rich domains in the outer leaflet of the PM via regulated exocytosis. (H) The relative ratio (percentage of above-threshold pixels, relative to all pixels in the thresholded image band) of D4 (red bars) and EGFP-D4 (green bars) signals in resting (REST) and stimulated (STIM) cells. Note the significant decrease in mCherry-D4-positive area and the significant increase in EGFP-D4 area after stimulation of regulated exocytosis, indicating that new cholesterol-rich domains are formed in the PM during the exocytotic fusion of vesicles with the PM. The numbers in the columns indicate the number of cells analyzed. Data are represented as means ± SEM. ***p < 0.001, **p < 0.01, *p < 0.05. Student’s t-test in (D) and (H); ANOVA with Holm-Sidak post-hoc test in (E) and (F).

Next, we assessed the degree of association between both markers. Insets (Figure 3C; arrows) indicate the types of associations between PRL- and D4-positive structures at the PM; some are overlapping or in contact (i.e., are associated) and others are separated (not associated). The distance between the non-associated structures was quantified (Figures S2C and S2D), revealing that depolarization-induced exocytosis significantly increased the D4-PRL distance (Figure 3D) from 0.28 ± 0.03 μm (REST, n = 182; nine cells) to 0.75 ± 0.13 μm (STIM, n = 96; ten cells, *p* < 0.001; see also Figure S2). Stimulation of exocytosis also significantly decreased the percentage of D4-PRL associations, as the overlap of D4- and PRL-positive structures at the PM decreased (Figure 3E) from 54.9% ± 2.0% in controls (REST, n = 9 cells) to 39.6% ± 2.5% in exocytosis-stimulated cells (STIM, n = 24 cells, *p* < 0.01). The decrease in the D4-PRL association in both experiments most likely reflects delivery of unlabeled vesicle-stored cholesterol to the PM during stimulated exocytosis.

To quantify the putative vesicle-stored cholesterol delivery, we modified the labeling protocol D by adding D4 to the exocytosis-stimulating solution (see Table S1, protocol E) to visualize the exocytosis-delivered cholesterol (Figure 3C [STIM + D4]). The degree of D4-PRL association was significantly higher in this group of cells (Figure 3E, red column, STIM + D4, 61.1% ± 3.2%, n = 16 cells) versus STIM (39.6% ± 2.5%, n = 24 cells, *p* < 0.01). Moreover, the D4-positive PM area (relative to the total PM area) was also significantly higher in the STIM + D4 group (Figure 3F; 9.9% ± 0.5%, n = 16 cells) versus the STIM group (7.7% ± 1.0%, n = 14 cells, *p* < 0.05).

To further test the delivery of vesicle cholesterol to the PM, the labeling protocols E and F (Table S1) were used, whereby the cholesterol-rich domains were first exposed to D4 (red fluorescence) to saturate the cholesterol-rich sites. Subsequently, we used a green fluorescent peptide, EGFP-D4, to visualize the newly added cholesterol after washing with vehicle (controls, REST) or stimulating exocytosis with 50 mM KCl (STIM, Figure 3G). In this experiment, the D4-positive pre-labeled (red) membrane area significantly decreased (Figure 3H) from 7.0% ± 0.5% in controls (REST, red bar, n = 20 cells) to 3.8% ± 1.6% in exocytosis-stimulated cells (STIM, red bar, n = 19 cells, *p* < 0.001). However, the green fluorescent EGFP-D4 PM area increased significantly from 1.2% ± 0.1% in controls (REST, green bar, n = 20 cells) to 1.6% ± 0.1 % in stimulated cells (STIM, green bar, n =19 cells, *p* < 0.01).

These data confirmed that vesicle cholesterol is delivered to the PM in the process of exocytosis, likely along the gradient between vesicle-stored and PM cholesterol.

### Exocytotic Release from Lactotrophs and Astrocytes is Stimulated by Cholesterol Extraction with MβCD

Exposing cells to 10 mM MβCD for 30 min reduced membrane cholesterol levels (Figure 2B) without affecting other membrane lipids (Table S2), consistent with previous reports (Churchward and Coorssen, 2009; Rituper et al., 2012). Next, we probed whether cholesterol extraction with MβCD affects exocytotic release in two physiologically distinct cell types: electrically excitable lactotrophs and electrically non-excitable astrocytes.

Application of MβCD to lactotrophs significantly increased the cumulative release of PRL over a period of 30 min (Figure 4A) from 0.54 ± 0.15 ng/mL in controls (REST) to 2.48 ± 0.10 ng/mL in treated cells (+MβCD, *p* < 0.01; n = 4 ELISA reactions). In astrocytes expressing ANP.emd peptide, which accumulates in secretory vesicles and can be released from cells by regulated exocytosis (Stenovec et al., 2014; Trkov et al., 2012), MβCD treatment (10 mM, 30 min) also increased the release of ANP.emd (Figure 4B), measured as changes in the bath fluorescence relative to the control. Assuming a linear relationship between the bath fluorescence intensity change and ANP.emd release, MβCD treatment triggered on average 26% more ANP.emd release compared with controls. MβCD treatment did not cause any significant cell death within the relevant time interval, indicating that cell lysis did not contribute to an increase in signal fluorescence.

**Figure 4.**
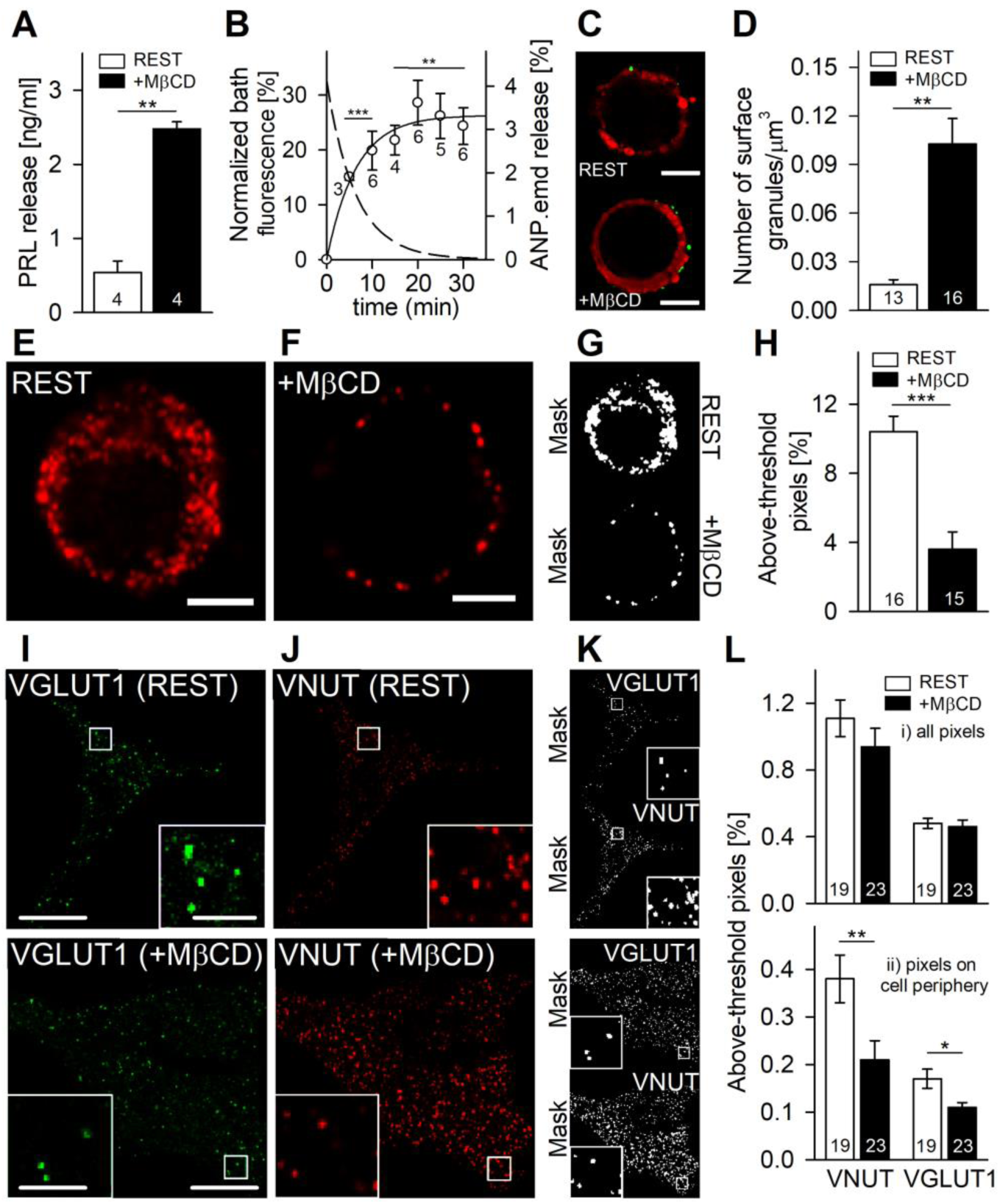
Acute Cholesterol Extraction with Methyl-β-cyclodextrin (MβCD) Stimulates Vesicle Cargo Release. (A) Cumulative release of prolactin (PRL) over 30 min from resting lactotrophs and those treated with 10 mM MβCD, measured with ELISA (the numbers in the columns represent the number of parallel experiments). (B) Normalized time-dependent increase in cell bath atrial natriuretic peptide (ANP.emd) fluorescence (%), relative to baseline (bath solution background); exponential fit (y = 26.31 × (1 − exp(−0.156t)), full line, t (time in min); and inverse function, y = 26.31 exp(–0.156t), dashed line, t (time in min) indicating time-dependent release of ANP.emd from MβCD-treated astrocytes (the numbers adjacent to the symbols represent the number of independent experiments). (C) Representative structured illumination microscopy images of resting (REST) and treated (+MβCD) lactotrophs (red, Vybrant DiD, a plasmalemma (PM) marker; green, surface-attached vesicle cargo-PRL granules). Note the increased surface presence of PRL granules after treatment with MβCD (+MβCD), indicating increased secretory activity. (D) Normalized number (to cell volume, μm^3^) of surface-attached PRL granules in resting (REST) and MβCD-treated (+MβCD) lactotrophs (see STAR Methods). (E and F) Representative confocal images of PRL vesicles (red) in (E) resting (REST) and (B) treated (+MβCD) lactotrophs. (G) Thresholded mask images of cells in (E) and (F). White, above-threshold pixels. Note the reduced number of PRL-positive vesicles in MβCD-treated lactotrophs (+MβCD). (H) The relative ratio of above-threshold PRL-positive pixels relative to all pixels in the cell region in resting (REST) and MβCD-treated (+MβCD) cells. (I) Representative confocal images of anti-VGLUT1-positive, i.e. glutamatergic vesicles (green), in resting (REST) and treated astrocytes (+MβCD). Insets display the enlarged framed areas; the bar depicts 2 μm. Note the reduced number of vesicles after treatment with MβCD. (J) Representative confocal images of anti-VNUT-positive, i.e. ATP-containing vesicles (red), in resting (REST) and treated astrocytes (+MβCD). Insets dsiplay the enlarged framed areas. Note the reduced number of vesicles after treatment with MβCD. (K) Thresholded mask images of cells in (I) and (J). White, above-threshold pixels. (L) The relative ratio of above-threshold pixels (%), relative to all pixels in (i) the entire cell region and in (ii) the cell region adjacent to the PM in resting (REST) and treated (+MβCD) astrocytes. The numbers in the columns indicate the number of cells analyzed. Data are presented as means ± SEM, ***p < 0.001, **p < 0.01, *p < 0.05. Student’s t test in (A), (C), (H), and (L); one-sample Student’s t-test versus 0 in (D). Scale bars, 3 μm in (B), (E) and (F); 10 μm (inset, 2 μm) in (I) and (J).

To validate these results at the single-cell level, we first imaged secreted PRL-containing granular content on the cell surface of lactotrophs (Cochilla et al., 1999; Mason et al., 1988) and the non-secreted, cytoplasmic, PRL-containing vesicles (Figures 4C–4F). During 30-min exposure to MβCD, the normalized number of PM surface-attached PRL granules significantly increased (Figure 4D) from 0.016 ± 0.003 granules/μm^3^ in controls (REST, n = 13 cells) to 0.103 ± 0.016 granules/µm^3^ in treated cells (+MβCD, n = 16 cells, *p* < 0.01). Similarly, exposure to MβCD decreased the quantity of cytoplasmic PRL-loaded vesicles in lactotrophs, indicating a substantial stimulation of vesicle cargo discharge during cholesterol extraction (Figure 4F). The surface area of above-threshold pixels (denoting cytoplasmic PRL vesicles; Figure 4G), relative to all pixels in the cell, significantly decreased (Figure 4H) from 10.4% ± 0.9% in controls (REST, n =16 cells) to 3.6% ± 1.0% in treated cells (+MβCD, n = 16 cells, *p* < 0.001).

MβCD treatment of astrocytes did not significantly change the above-threshold vesicular nucleotide transporter (VNUT) signal that represent ATP-containing vesicles (Gundersen et al., 2015) (Figure 4L, top graph), nor the above-threshold vesicle glutamate transporter 1 (VGLUT1) signal that represents glutamate-containing vesicles (Stenovec et al., 2008) (Figure 4L, top graph). However, we observed significantly less VNUT- and VGLUT1-positive vesicles in the 2-µm-thick band region, adjacent to the PM (Figure 4L, bottom graph). In this region, MβCD treatment significantly decreased the level of above-threshold VNUT-positive pixels from 0.38% ± 0.05% in controls (VNUT, REST, n=19 cells) to 0.21% ± 0.04% in treated cells (VNUT, +MβCD, n = 23 cells, *p* < 0.01) as well as the amount of above-threshold VGLUT1-positive pixels from 0.17% ± 0.02% in controls (VGLUT1, REST, n = 19 cells) to 0.11% ± 0.01% in treated cells (VGLUT1, +MβCD, n = 23 cells, *p* < 0.05).

These results demonstrate that MβCD-induced cholesterol extraction facilitates exocytotic release from both of the cell types studied.

### The Cholesterol Extraction-Mediated Increase in Vesicle Release is Independent of Changes in Cytosolic Free Ca^2+^ Levels

It is conceivable that MβCD cell treatment enhances the observed exocytotic release through an unspecific increase in cytosolic free Ca^2+^ concentration ([Ca^2+^]_i_), a stimulus for regulated exocytosis in both cell types (Kreft et al., 2004; Krzan et al., 2003; Zorec et al., 1991). Therefore, we measured the changes in [Ca^2+^]_i_ in lactotrophs and astrocytes during the exposure to 10 mM MβCD or 10 mM cholesterol-replenishing solution. During MβCD treatment [Ca^2+^]_i_ increased in lactotrophs but decreased in astrocytes (Figure S4). Therefore, these results indicate that cholesterol depletion may promote the discharge of vesicle in a [Ca^2+^]_i_-independent manner by affecting the fusion pore diameter.

### A Theoretical Description of Cholesterol-Mediated Fusion Pore Properties

We developed a model that describes the mechanism of cholesterol redistribution between the vesicle and the PM along a cholesterol gradient, passing through the highly curved membranous fusion pore (Figures 5A and 5B). At the fusion pore orifice, a shift in the membrane curvature occurs, from a negative curvature in the luminal face of the vesicle membrane to a positive one at the fusion pore orifice. The change in the curvature affects the properties of the PM (Yesylevskyy et al., 2017), forming a boundary for membrane molecules with negative spontaneous curvature, such as cholesterol, to be transferred from the vesicle membrane to the PM. This generates tension (Baumgart et al., 2003; Julicher and Lipowsky, 1996) and creates a radial force, constricting the fusion pore diameter (Figure 5C). As the fusion pore diameter at steady state depends on the vesicle diameter (Flasker et al., 2013; Jorgacevski et al., 2010; Shin et al., 2018), this model also considers this relationship (Figure 5D). Details of how this model was developed are provided in the Supplemental Information (Figure S6).

**Figure 5.**
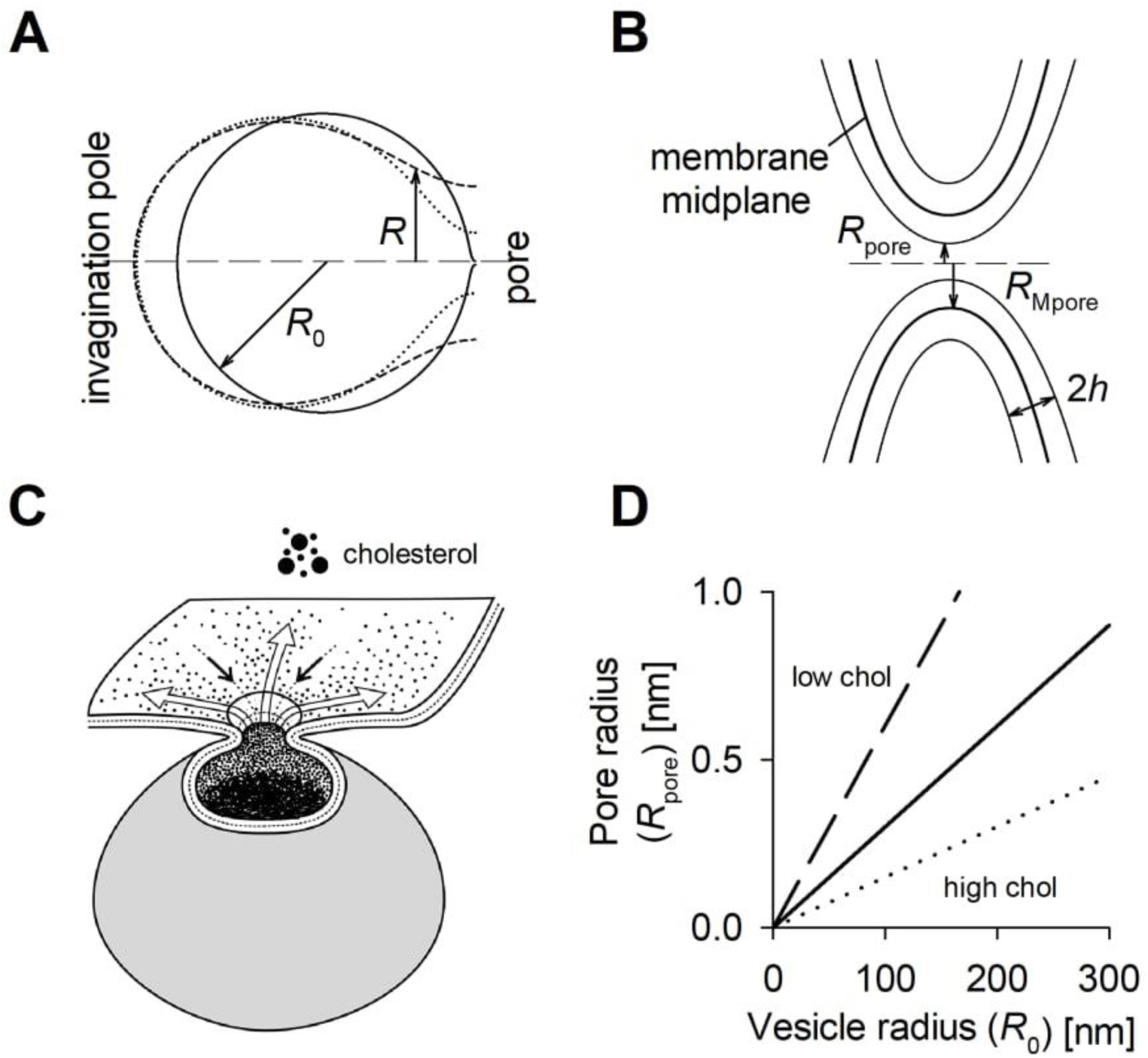
A Model of the Fusion Pore Radius as a Function of Vesicle Radius at Different Cholesterol Concentrations between the Vesicle and Plasmalemma. (A) Longitudinal cross-sections of characteristic invagination shapes (secretory vesicle connected by the fusion pore to the PM) at a given invagination area 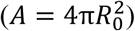 for *R*_0_*c*_p =_ 1.003 (full line), 1.045 (dotted line), and 1.099 (dashed line); *R*_0_ denotes the vesicle radius, *R* denotes the distance of the vesicle membrane from the symmetry axis. (B) Scheme of the fusion pore, where *R*_pore_ denotes the fusion pore radius and *R*_Mpore_ is the radius of the membrane midplane at the fusion pore. (C) A scheme of the fusion pore and processes taking place when the fusion pore opens. Vesicle cholesterol density is depicted to be very high in the vesicle membrane as well as in the vesicle lumen, likely part of a crystalline array with hormones. Cholesterol in the intraluminal, a negatively curved face of the vesicle membrane, exits the vesicle through the fusion pore. However, the positive membrane curvature at the fusion pore orifice represents a boundary/barrier for cholesterol exiting the vesicle into the outer leaflet of the PM. The white arrows denote the direction of cholesterol transfer from the vesicle toward the outer leaflet of the PM. This creates tension and a radial force (tips of the black arrows), keeping the fusion pore radius constricted. Cholesterol molecules transferred from the vesicle membrane mix with cholesterol-rich rafts in the outer-leaflet of the PM.. See Supplemental Information for the development of the model. (D) Dependence of the fusion pore radius (*R*_pore_) as a function of the vesicle radius (*R*_0_) for the ratio β/*E*_0 =_ 6 × 10^−3^ (dashed line), 3 × 10^−3^ (full line), and 1.5 × 10^−3^ (dotted line) (see Supplemental Information, eqs. 6 and 7, where constants *E*_0_, β, and α are defined) at relatively large α, representing low, normal, and high cholesterol (chol.) in the PM, respectively.

In brief, line tension at the fusion pore, creating a radial force constricting the fusion pore diameter (Figure 5C; Supplemental Information, eq. 5), is proportional to the difference in cholesterol concentrations between the vesicle membrane and the PM (Supplemental Information, eq. 6). Due to its anisotropic and negative spontaneous curvature, cholesterol determines the bending elasticity of the membrane and preferentially accumulates in membrane domains with relatively high local negative curvature (Chen and Rand, 1997; Jorgacevski et al., 2010). Therefore, the cholesterol density in the membrane is larger at more negatively curved membranes (Yesylevskyy et al., 2017), i.e. at smaller fusion pore radii with respect to the secretory vesicle size (Supplemental Information, eq. 7). By simulating the influence of the difference in cholesterol concentrations between the vesicle membrane and the PM, we found a linear dependence of fusion pore radius on vesicle radius (Figure 5D; Supplemental Information, eq. 9), where the slope of this relationship is determined by cholesterol-related parameters (Supplemental Information). Increased versus reduced cholesterol content is predicted to result in a relatively narrow versus wide fusion pore radius, respectively (Figure 5D).

### Fusion Pore Conductance Increases in Cholesterol-Depleted Lactotrophs and Astrocytes

To verify the role of cholesterol in determining the vesicle fusion pore properties (Figure 5), unitary exocytotic events were monitored by high-resolution membrane capacitance (C_m_) measurements in lactotrophs and astrocytes under control, cholesterol-depleted, and cholesterol-replenished conditions (Table S3).

In controls, ∼64% of patch-clamped lactotrophs (52 of 81) exhibited upward steps in C_m_, and 95% (n = 1354) of those were reversible, meaning that a discrete upward step in C_m_ was followed by a downward step of similar amplitude within 3 s (Gucek et al., 2016) (Figure 6A). Similar reversible events were observed in astrocytes (Figure 6B). These consecutive steps in C_m_ reflect transient exocytotic fusion pore openings (Alvarez de Toledo et al., 1993; Zorec et al., 1991), whereas irreversible upward steps in C_m_ (i.e. full fusion, Figure 6A) represent a complete merger of the vesicle membrane with the PM (Neher and Marty, 1982). Similar to a previous study (Rituper et al., 2012), the 10 mM MβCD pretreatment (Table S2) decreased the frequency of reversible fusion pore opening events (Figure 6C, Table S3). In cholesterol-replenished cells, the frequency of reversible fusion pore events was significantly increased, whereas full fusion events decreased. Moreover, the frequency of irreversible events was not affected by cholesterol depletion in lactotrophs; however, after cholesterol replenishment, the frequency of irreversible events decreased slightly (Table S3).

**Figure 6.**
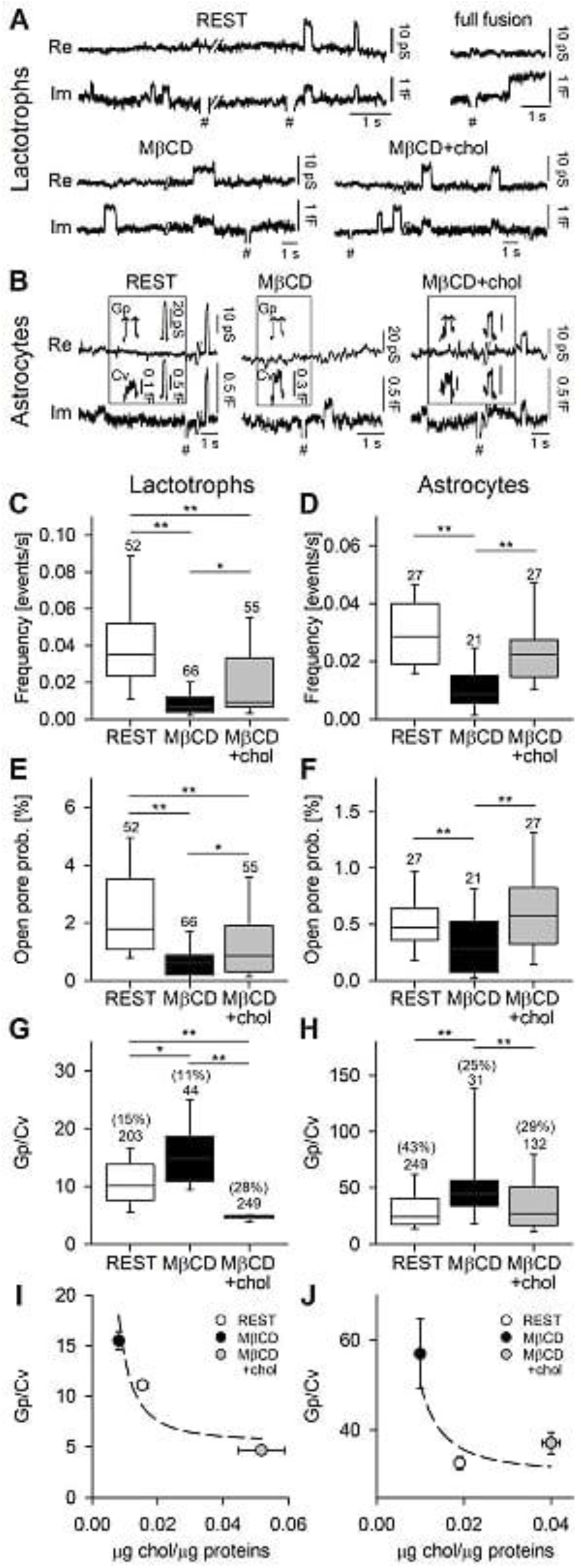
Lowering Cholesterol Decreases the Probability of Vesicle Fusion and Increases Fusion Pore Conductance in Lactotrophs and Astrocytes, an Effect Rescued by Cholesterol Replenishment. (A) Representative patch-clamp cell-attached membrane recordings of Re (real) and Im (imaginary) parts of admittance traces displaying reversible and irreversible Im discrete steps (proportional to the membrane capacitance, C_m_) with or without projections to Re, in resting (REST), cholesterol-extracted (+MβCD), and cholesterol-replenished (MβCD + chol) lactotrophs, after MβCD washout. (B) Representative Re and Im traces displaying reversible discrete Im steps with or without projections to Re in resting (REST), (+MβCD), and (MβCD + chol) treated astrocytes. Cells in the +MβCD group in (A) and (B) were pretreated with MβCD (10 mM, 30 min) before recordings. After cholesterol extraction and before recording, cholesterol was replenished (MβCD + Chol) with cholesterol-replenishing solution (10 mM, 30 min). (C and D) The frequency of reversible fusion events in resting (REST), (+MβCD), and (MβCD + chol) lactotrophs (C) and astrocytes (D). (E and F) Open (detectable) fusion pore probability in resting (REST), (+MβCD), and (MβCD + chol) lactotrophs (E) and astrocytes (F). (G and H) Fusion pore conductance (G_p_) normalized to vesicle membrane capacitance (C_v_) (G_p_/C_v_) in resting (REST), (+MβCD), and (MβCD + chol) treated lactotrophs (G) and astrocytes (H). The numbers in parentheses denote the percentage of Re projections to the Im trace, relative to all transient Im events. (I-J) Normalized fusion pore conductance (G_p_/C_v_), proportional to (*R*_pore_/*R*_*0*_)^2^ (Supplemental Information, eq. 7) versus normalized cholesterol concentration (µg chol/µg protein), proportional to *E*_*1*_ (Supplemental Information) in resting (REST), (+MβCD), and (MβCD + chol) (I) lactotrophs and (J) astrocytes. Fitted (to all data pairs) inverse square curve is of the form (G_p_/C_v_) = (8.23×10^−4^ ± 4.35×10^−5^)/x^2^) + (5.33 ± 0.23), r = 0.649 (I) and. (G_p_/C_v_) = (2.00×10^−3^ ± 5.56×10^−4^)/x^2^) + (30.59 ± 1.95), r = 0.174 (J). The numbers above the box plots indicate the number of cells analyzed in (C)–(F) and the number of transient Im events with Re projections in (G) and (H). Data are presented as medians with interquartile range (C–H) and means ± SEM in (I and J). **p < 0.01, *p < 0.05. ANOVA on ranks with Dunn’s post-hoc test. # denotes 10 fF calibration.

The probability of observing a fusion pore in the open state, calculated as the sum of all dwell times of reversible C_m_ events divided by the time of the recording, decreased significantly after cholesterol depletion (*p* < 0.01;Figure 6E) and was partially, yet significantly, restored after cholesterol replenishment to 0.86% (*p* < 0.01 versus control; Table S3). The fusion pore conductance (G_p_), a measure of the fusion pore diameter, can be calculated for transient, i.e. reversible C_m_ events that display a projection from the imaginary (Im) to the real (Re) part of the admittance signal (Lollike et al., 1995). Such C_m_ events comprised 15% of all events in control (203 of 1354 events), 11% in cholesterol-depleted cells (44 of 400 events), and 28% in cholesterol-replenished cells (249 of 877 events; Figure 6G). The settings of our experimental system allowed the detection of fairly narrow diameter fusion pores (0.3–3 nm) (Jorgacevski et al., 2010); therefore, the decrease in the relative occurrence of projected C_m_ events in MβCD-treated cells indicated that cholesterol extraction promoted the formation of fusion pores with wider diameters, which were no longer detectable as a projection between the Im and Re parts of the admittance signals. Consistent with this, the G_p_/C_v_ ratio, in which the fusion pore conductance (G_p_) is normalized to the vesicle size (C_v_), significantly increased in cholesterol-depleted cells (Figure 6G, p < 0.05; Table S3). In cholesterol-replenished lactotrophs, the G_p_/C_v_ ratio decreased significantly in comparison with controls (*p* < 0.01; Table S3). The cholesterol-mediated changes in G_p_/C_v_ correlated with the measured cholesterol concentration (Table S2 and Figure 6I).

Similar results were obtained in astrocytes. The frequency of reversible C_m_ events significantly (*p* < 0.01) decreased in MβCD-treated cells and was restored (*p* < 0.01) in cholesterol-replenished cells (Figure 6D). Unlike in lactotrophs, we observed a significant decrease in the frequency of irreversible events in cholesterol-depleted astrocytes, which was restored by cholesterol replenishment. The percentage of reversible events, in comparison with all events, decreased in cholesterol-depleted astrocytes and was restored by cholesterol replenishment (Table S3).

The probability of observing the fusion pore open state (Figure 6F) was significantly reduced by cholesterol depletion (*p* < 0.01; Table S3). Cholesterol replenishment fully restored the open fusion pore probability (*p* < 0.01 versus cholesterol-depleted cells; Table S3). Similar to recordings in lactotrophs, there was a decrease of reversible events with projections (representing relatively narrow fusion pore diameters) from the Im to Re parts of the admittance (Lollike et al., 1995), from 43% (249 of 576 events) in controls to 25% (31 of 122 events) in cholesterol-depleted astrocytes; this increased to 29% (132 of 462 events) with cholesterol replenishment. Accordingly, the G_p_/C_v_ ratio (Figure 6H) in astrocytes increased significantly after cholesterol depletion (*p* < 0.01) and was fully restored by cholesterol replenishment (*p* < 0.01 versus cholesterol-depleted cells; Table S3). The cholesterol-dependent G_p_/C_v_ correlated with the measured cholesterol (Figure 6J).

These results demonstrate that relatively high cholesterol levels constrict fusion pores, whereas cholesterol depletion widens fusion pores, consistent with our theoretical predictions (Figure 5D) as further established in Figure S5, which display the relationship between the fusion pore diameter (estimated from G_p_) and the vesicle diameter (determined from C_v_) (lines fitted to eq. 9: Supplemental Information).

### Reduced Fusion Pore Conductance (G_p_) in Cholesterol-Enriched Vesicles of Niemann-Pick Disease Type C1 Null Mouse Fibroblasts

The fibroblast cell model of lysosomal storage disease, in which the Npc1 protein, the vesicle cholesterol transporter, is deleted, demonstrates an accumulation of cholesterol in the vesicle lumen and membrane (Liscum, 2000; Mukherjee and Maxfield, 2004). Confocal microscopy of fixed cultured cells revealed abundant D4-positive vesicles in the cytoplasm of *Npc1*^−*/*−^ cells (Figures 7A and 7B). The number of D4-positive vesicles (per cell plane) was ∼2.7 fold higher in *Npc1*^−*/*−^ cells (*p* < 0.01, n = 15) compared with wild-type fibroblasts (n = 15; Figure 7C), and the cumulative fluorescence intensity of D4-positive vesicles was ∼2.6-fold higher in *Npc1*^−*/*−^ (*p* < 0.001) than in wild-type fibroblasts (Figure 7D). These data indicate that fibroblasts lacking functional Npc1 accumulate cholesterol in vesicles, presumably in late endosome/lysosomes (Mukherjee and Maxfield, 2004), consistent with a previous study (Higgins et al., 1999).

**Figure 7.**
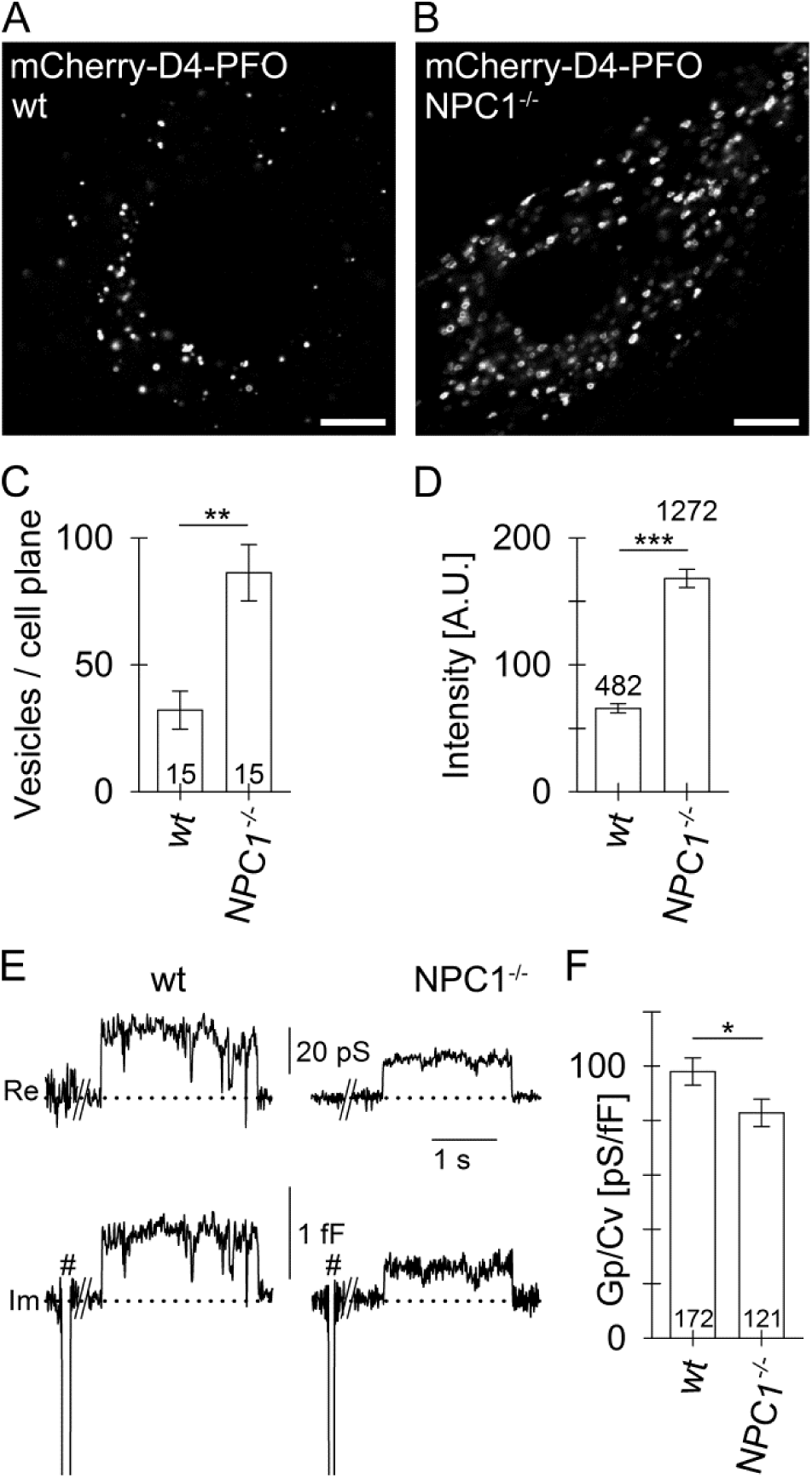
Increased Cholesterol Content in Fibroblasts Lacking Npc1 is Associated with Reduced Fusion Pore Conductance. (A and B) Confocal images of fixed wild-type (wt) and *Npc1*^−*/*−^ fibroblasts labeled with the cholesterol-specific membrane-binding domain of perfringolysin O (mCherry-D4) reveal numerous vesicle-like structures in the *Npc1*^−*/*−^ versus wt cells (scale bar, 10 μm). (C and D) Number (mean ± SEM) of mCherry-D4-positive vesicles per cell plane (C) and the cumulative vesicle fluorescence intensity (D) in wt and *Npc1*^−*/*−^ fibroblasts. The numbers at the bottom and top of the bars indicate the number of cell images and vesicles analyzed, respectively; **p < 0.01, ***p < 0.001 versus respective comparisons (Mann-Whitney U test). (E) Electrophysiological recordings of real (Re, top) and imaginary (Im, bottom) parts of the admittance signals in vesicles transiently interacting with the PM in wt and *Npc1*^−*/*−^ fibroblasts. Dotted lines indicate baseline level; # denote calibration pulses. (F) Fusion pore conductance (G_p_) normalized to vesicle capacitance (C_v_, G_p_/C_v_) is reduced in *Npc1*^−*/*−^ fibroblasts. The numbers at the bottom of the bars indicate the number of vesicles analyzed. *p < 0.05 (Mann-Whitney U test).

High-resolution C_m_ measurements revealed that G_p_, normalized to vesicle capacitance (G_p_/C_v_), decreased from 98 ± 5 pS/fF in controls (n = 172) to 83 ± 5 pS/fF in *Npc1*^−*/*−^ fibroblasts (n = 121, *p* < 0.05, Figure 7F), indicating that increased vesicle cholesterol in *Npc1*^−*/*−^ fibroblasts narrows the fusion pore.

## DISCUSSION

Several stages of exocytosis were proposed (Rand and Parsegian, 1986) beginning with the apposition of membranes destined to fuse, which is then ending by (1) destabilization of the membranes in contact, (2) coalescence of the two membrane surfaces and (3) re-stabilization of the fused membranes. These end stages were considered to occur within microseconds to maintain the cell’s integrity. Hence, this view prompted the idea that once the fusion pore is formed, the fusion pore must widen abruptly (full-fusion exocytosis). However, more recent experimental evidence revealed that the fusion pore, once established, enters a stable, dynamically regulated state, with diameters ranging between sub-nanometers and several hundred nanometers (Shin et al., 2018; Vardjan et al., 2007). These pores can then undergo either reversible constriction or closure to limit vesicle discharge or full widening (full-fusion exocytosis) to boost vesicle discharge (Kreft et al., 2018). It is generally acknowledged that these transitions are regulated by proteins (Shin et al., 2018), which may also form the fusion pore walls (Almers and Tse, 1990; Chang et al., 2017). Although the fusion pore was also modeled to be exclusively lined by lipids (Nanavati et al., 1992), it is more likely to be a mixture of proteolipids (Gasman and Vitale, 2017; Sharma and Lindau, 2018). There has been interest in how cholesterol initiates and regulates the stages of regulated exocytosis (Churchward et al., 2005a; Cookson et al., 2013; Mahadeo et al., 2015; Najafinobar et al., 2016; Rituper et al., 2012; Rogasevskaia and Coorssen, 2006; Stratton et al., 2016; Wang et al., 2010; Xu et al., 2017); however, direct evidence that membrane lipids define the dynamics of single fusion pore stages is missing. Our results establish the role of a cholesterol-dependent radial force that constricts the fusion pore (Figures 5 and 6). This lipid-based mechanism of fusion pore regulation provides the missing evidence proving that the fusion pore itself functions as a proteolipidic entity.

Four lines of evidence support this conclusion. First, the spontaneous negative curvature of cholesterol, a structural stress factor (Rand and Parsegian, 1986), likely promotes the first step of the membrane merger (Churchward and Coorssen, 2009). However, cholesterol may also play a role during post-fusion stages. Upon stimulating regulated exocytosis, the fusion pore is formed, through which vesicle cholesterol exits and subsequently mixes with the outer leaflet of the PM, contributing to an asymmetric composition of the PM. Cholesterol has been shown to be unevenly distributed in various cellular membranes (Ikonen, 2018). In some cell types, cholesterol accumulates in lysosomes, especially in diseased states (Liscum, 2000; Mukherjee and Maxfield, 2004), and in the vesicle membranes of secretory cells (Bogan et al., 2012; Mahadeo et al., 2015). Consistent with these previous studies, we found that PRL-storing vesicles are rich in cholesterol (Figure 1). PRL is discharged from vesicles through regulated exocytosis (Akerman et al., 1991; Stenovec et al., 2004), and it appears that vesicle cholesterol is transferred to the PM along with the hormone. Stimulation of regulated exocytosis reduced the density and size of the mCherry-D4-PFO (D4) labeled cholesterol-rich PM domains (Figure 2). This likely reflects the delivery of vesicle cholesterol together with the hormone discharge, as the surface PM deposits of fluorescently labeled hormone from vesicles correlated with increased outer leaflet PM cholesterol domains (Figure 3). Both hormones and cholesterol exit vesicles along a gradient; hormones are released into the extracellular space and cholesterol into the PM. Furthermore, cholesterol exiting the vesicle through the fusion pore may also regulate it, as tested in subsequent experiments.

The second line of evidence is that cholesterol depletion widens the fusion pore independently of [Ca^2+^]_i_. Exposing cells to MβCD while monitoring amperometric discharge of vesicle cargo (Cookson et al., 2013; Wang et al., 2010) revealed that cholesterol enrichment increased the persistence of the pre-foot spike of the amperometric signal, which is considered to indirectly represent the leakage of vesicle cargo through a narrow fusion pore before the pore dilates for full exocytotic release (Chow et al., 1992). We monitored vesicle discharge in electrically excitable pituitary lactotrophs and in electrically non-excitable astrocytes. Vesicle cargo, labeled by immunofluorescence in lactotrophs or by the expression of fluorescently tagged peptide in astrocytes, was secreted with MβCD treatment (Figure 4). Direct monitoring of fusion pore conductance, a measure of fusion pore diameter, revealed that cholesterol depletion widens and cholesterol enrichment narrows the fusion pore in both cell types (Figure 6).

Perturbations of membrane cholesterol may affect the activation of several types of Ca^2+^ channels, including voltage-gated Ca^2+^ channels, thus affecting [Ca^2+^]_i_ (Levitan et al., 2010). Therefore, MβCD may stimulate regulated exocytosis indirectly (Wang et al., 2010) through an increase in [Ca^2+^]_i_. In contrast, whereas an MβCD-dependent increase in [Ca^2+^]_i_ was recorded in lactotrophs, a prominent reduction was recorded in astrocytes (Figure S3), indicating that MβCD-dependent cholesterol depletion in astrocytes widens the fusion pore in the absence of an increase in [Ca^2+^]_i_, and that fusion pore widening is due to direct, cholesterol-driven regulation of the fusion pore.

Third, the molecular shape and physicochemical properties of cholesterol determine membrane processes through effects on membrane curvature, including the initial membrane merger step of regulated exocytosis (Churchward et al., 2008b; Jorgacevski et al., 2010; Rand and Parsegian, 1986). Arguably, membrane lipids segregate to different parts of cellular membranes based on their intrinsic curvature (Mukherjee et al., 1999) in addition to other mechanisms of lipid segregation (Tian and Baumgart, 2009). Cholesterol distributes unevenly in cellular compartments (Ikonen, 2018; Wustner and Solanko, 2015). The walls of the open fusion pore represent a highly curved structure in contact with the PM. This orifice represents a boundary across which the cholesterol gradient between the vesicle and the PM exerts tension. Due to this tension, a radial force is generated, which prevents fusion pore widening (Figure 5C). An analogy for this are the capillary forces at the boundary of a water-filled glass, which increase the water level at the glass surface. The radial force at the orifice of the fusion pore is assumed to depend on the difference in cholesterol concentration between the vesicle and the PM, driving the redistribution of vesicle cholesterol into the outer leaflet of the PM (Figure 2). Based on this, a theoretical model of how cholesterol controls the fusion pore diameter was developed (Supplemental Information). This model takes into account that the vesicle diameter determines the fusion pore radius, as confirmed experimentally (Flasker et al., 2013; Jorgacevski et al., 2010; Shin et al., 2018). Our model has multiple solutions for a stable fusion pore radius, in which the relative interaction is proportional to the interaction energy per surface area (Supplemental Information), and is, depending on the highly curved membranes at the fusion pore (Figure 5B), plotted as a function of the normalized fusion pore radius (marked by arrows in Figure S6A). This makes this model suitable for describing not only very narrow fusion pores (Vardjan et al., 2007), but also those with much larger diameters (Shin et al., 2018). The radial force defined in this model depends linearly on the local cholesterol availability (eq. 6; see also the direction of the forces depicted in Figure 5C). We estimated the force at the fusion pore (γ_pore_) from the dimensionless transfer shear force (Γ_pore_) in the radial direction: Γ_pore =_ *R*_0_γ_pore_/8π*k*_*c*_, where *R*_*0*_ is the vesicle radius and *kc* is the bending modulus (Supplemental Information). For a typical value of Γ_pore_ = 0.4 (Figure S6B), using *R*_*0*_ = 100 nm and *k*_*c*_ = 20k_B_T (Rawicz et al., 2000), the estimate for γ_pore_ is 80 pN, essentially the same as the value reported recently for the fusion pore in chromaffin cells (Shin et al., 2020). A similar range of forces affecting the fusion pore can be predicted by considering the interactions between SNARE proteins at post-fusion stages of exocytosis (Jorgacevski et al., 2011).

The validity of this model is corroborated by the series of experiments (Figures 6I and 6J), in which normalized fusion pore conductance (G_p_/C_v_) is correlated with cholesterol content, indicating that changes in cellular cholesterol content may regulate the vesicle fusion pore in health and disease.

Fourth, our results provide evidence supporting the role of cholesterol redistribution between the vesicle and PM in the regulation of the exocytotic fusion pore, as disease-related increases in vesicle cholesterol affected the fusion pore (Figure 7F) in a manner consistent with predictions by the model (Figure 5D). In Niemann-Pick disease type C1 mutations in NPC1, a protein involved in lysosome: ER contact site formation and therefore indirectly facilitating sterol transport (Hoglinger et al., 2019; Winkler et al., 2019), lead to an accumulation of vesicle cholesterol (Liscum, 2000; Mukherjee and Maxfield, 2004). In *Npc1*^−*/*−^ fibroblasts vesicles contained on average ∼2-fold more cholesterol (Figure 7A-D). High-resolution membrane capacitance measurements of fusion pore conductance, normalized to vesicle capacitance (G_p_/C_v_), revealed fusion pore constriction by over 15% in these fibroblasts (Figure 7). These data demonstrate that the disease-related increase in vesicle cholesterol in Npc1−/− fibroblasts narrows the fusion pore, which arguably hinders the capacity of vesicle cargo release. The contribution this makes to the pathophysiology of NPC therefore merits further investigation.

Moreover, it is possible that pharmacologically induced changes in the PM cholesterol levels may also affect the fusion pore and therefore indirectly affect cell function(s). Recently, it has been demonstrated that ketamine alters the density of cholesterol in the PM (Lasic et al., 2019). This anesthetic and psychotomimetic has gained much attention due to its rapid and long-lasting antidepressant effects (Berman et al., 2000). Ketamine was previously considered a non-competitive N-methyl-D-aspartate receptor antagonist (Sanacora et al., 2012), with an antidepressant action owing to increased synthesis of brain-derived neurotrophic factor (BDNF) (Li et al., 2010), However ketamine has now also been shown to inhibit the release of BDNF from astrocytes, likely by reducing [Ca^2+^]_i_ (Stenovec et al., 2016) and stabilizing the fusion pore in a narrow flickering configuration (Lasic et al., 2016). These effects are very likely due to the relatively rapid, ketamine-invoked changes in cholesterol density in the PM (Lasic et al., 2019), indicating that drug-induced changes in PM cholesterol may shape the signaling capacity of cells through regulation of the fusion pore

In summary, this paper demonstrates that stimulation of exocytosis leads to the transfer and subsequent mixing of vesicle cholesterol with the outer leaflet of the PM. Cholesterol redistribution through the fusion pore instigates constriction of the fusion pore through a radial force, depending on the difference in cholesterol concentrations between the vesicle and the PM. This theory provides a new foundation for understanding this ubiquitous membrane intermediate of eukaryotic cells, not only in exocytosis but also in endocytosis, intracellular trafficking, fertilization, and viral membrane penetration, for which membrane pore dynamics is a key process that is now clearly shown to involve both membrane proteins and lipids.

## Supporting information

Supplemental info on Cholesterol-mediated fusion pore regulation

## ACKNOWLEDGMENTS

This work was supported by grants from the Slovenian Research Agency (P3 310, P1-0055, J3 4051, J3 4146, L3 3654; J3 3236, J3 6790, J3 6789, J3 7605), CIPKEBIP, COST Nanonet, COST Mouse Ageing, COST CM1207 – GLISTEN. JRC acknowledges the support of the NHMRC (Australia; APP1065328) and Brock University. F.M.P. is a Royal Society Wolfson Research Merit Award holder and a Wellcome Trust Investigator in Science. Claire Smith from the Platt Lab is acknowledged for the technical expertise with the *Npc1* knockout and control fibroblasts..

## AUTHOR CONTRIBUTIONS

B.R., A.G., M.L., U.G., S.T.B., E.L., M.B. and M.S. carried out electrophysiology and imaging experiments and performed data analysis. P.S.A. and J.R.C. carried out the lipidomic analysis after cholesterol depletion in cells. A.Š. and G.A. developed and prepared the D4 cholesterol fluorescent marker. F.M.P. provided the *Npc1* deficient and control fibroblasts. B.B. developed the cholesterol-mediated radial force membrane fusion pore constriction model. R.Z., J.R.C., B.B., J.J., A.V., M.K., and M.S. discussed the design of the experiments. R.Z. directed and wrote the manuscript with help from all authors.

## SUPPLEMENTAL FIGURES

**Figure S1.**
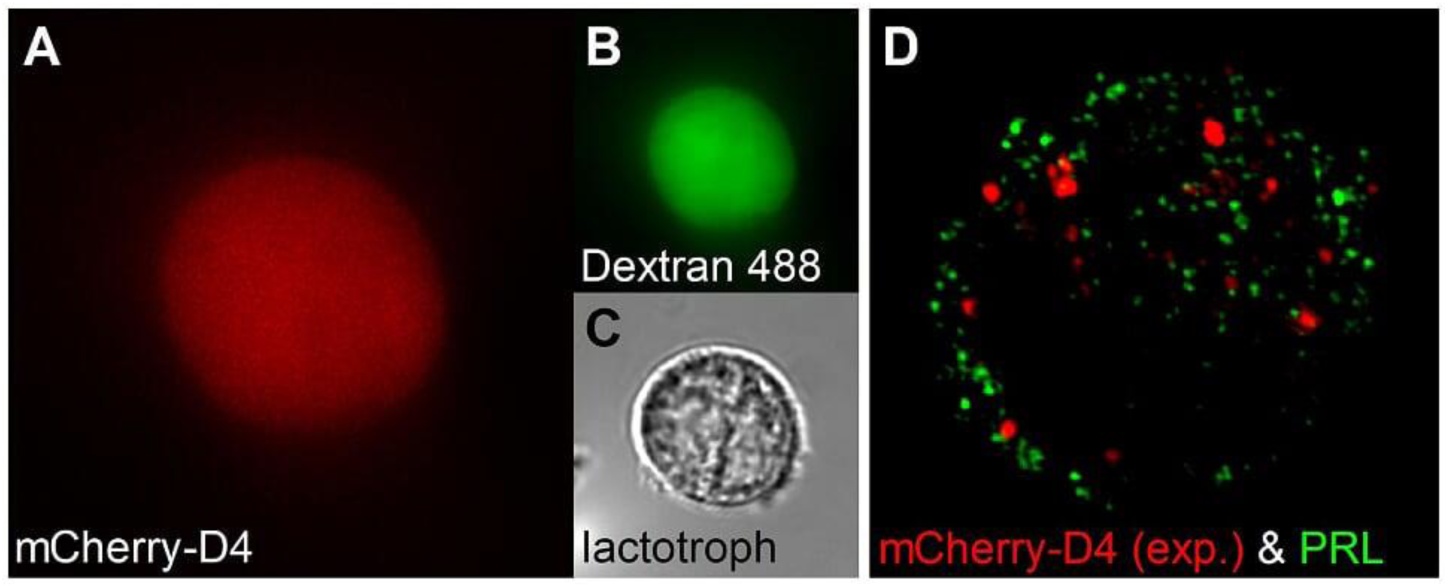
Microinjected mCherry-D4-PFO Poorly Labels the Cytoplasmic Leaflet of the Plasmalemma and the Membrane of Prolactin (PRL) Vesicles, Related to Figure 2. (A–C) Representative fluorescent and transmission images of a lactotroph microinjected with mCherry-D4-PFO or mCherry-D4-PFO-YQDA (red, A) alongside with Dextran Alexa 488 (green, B). (C) Transmission image of the cell shown in A & B. (D) Representative super-resolution microscopy image of a lactotroph. Red, expressed plasmid mCherry-D4-PFO by electroporation (mCherry-D4 (exp.), see STAR Methods); green, anti-prolactin antibody labeling, PRL).

**Figure S2.**
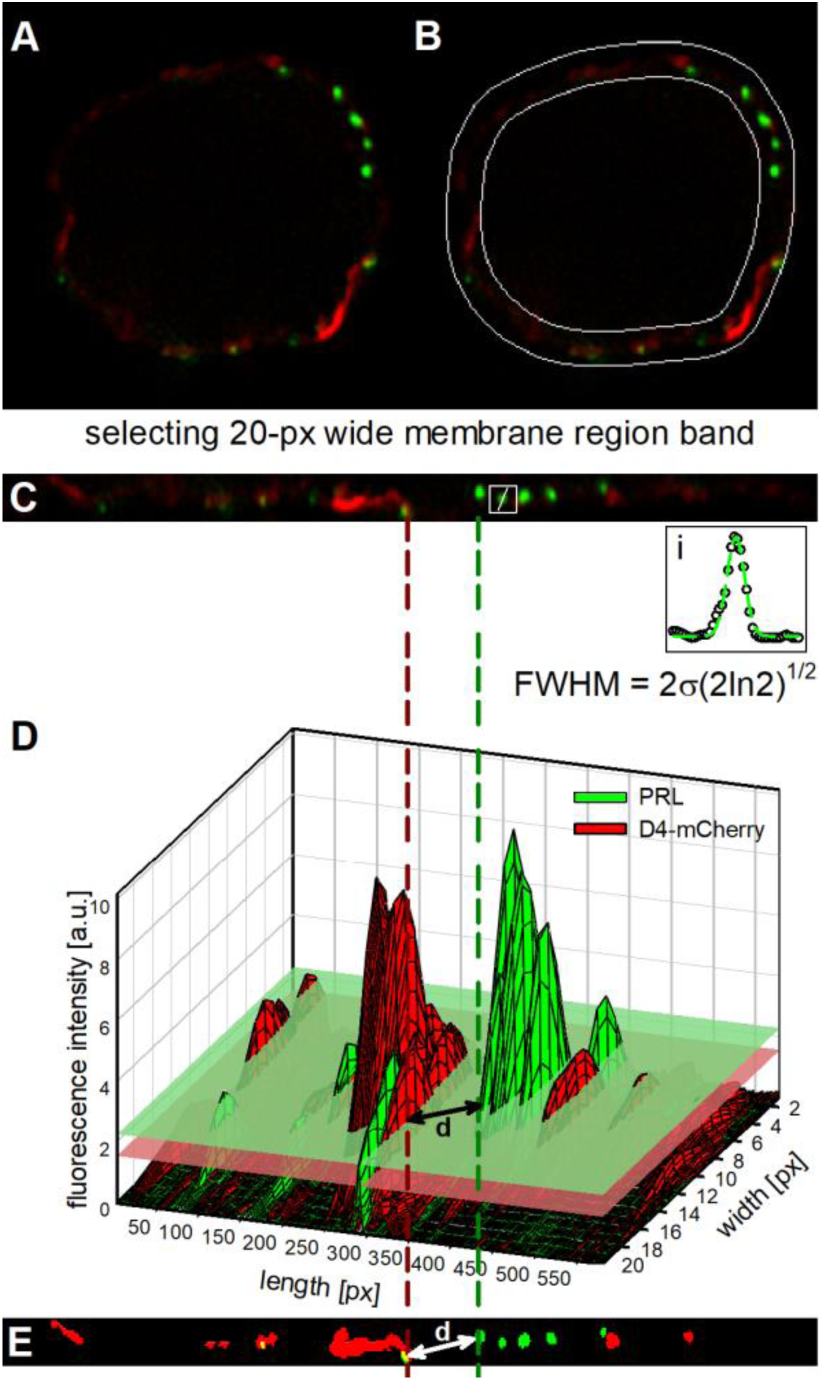
Graphical Representation of mCherry-D4-PFO (D4) and the Prolactin (PRL) Analysis Procedure, Related to Figure 2. (A) Representative structured illumination microscopy image of a lactotroph labeled with mCherry-D4-PFO (red) and PRL (green). (B) A band of the peri-plasmalemmal content of fluorescent markers. White, 20-pixel (∼800 nm)-wide band encompassing the plasmalemmal region of the cell in (A). (C) Straightened (linear) 20-pixel-wide structured illumination microscopy image of the plasma membrane area. Inset (i): a graphical representation of the vesicle size calculation method (vesicle diameter = FWHM = 2σ√(2ln2); σ is the standard deviation of the Gaussian fit). (D) Graphical-3D representation of the thresholding protocol. Red D4 signal; green PRL signal; red plane, D4 threshold; green plane, PRL threshold (see also STAR Methods). Px-pixels for length and width. (E) Thresholded 20-pixel-wide image of the plasma membrane region of the cell in (D). Red, above-threshold D4 signal; green, above-threshold PRL signal. d is the shortest distance from the D4 domain to the PRL-labeled structure.

**Figure S3.**
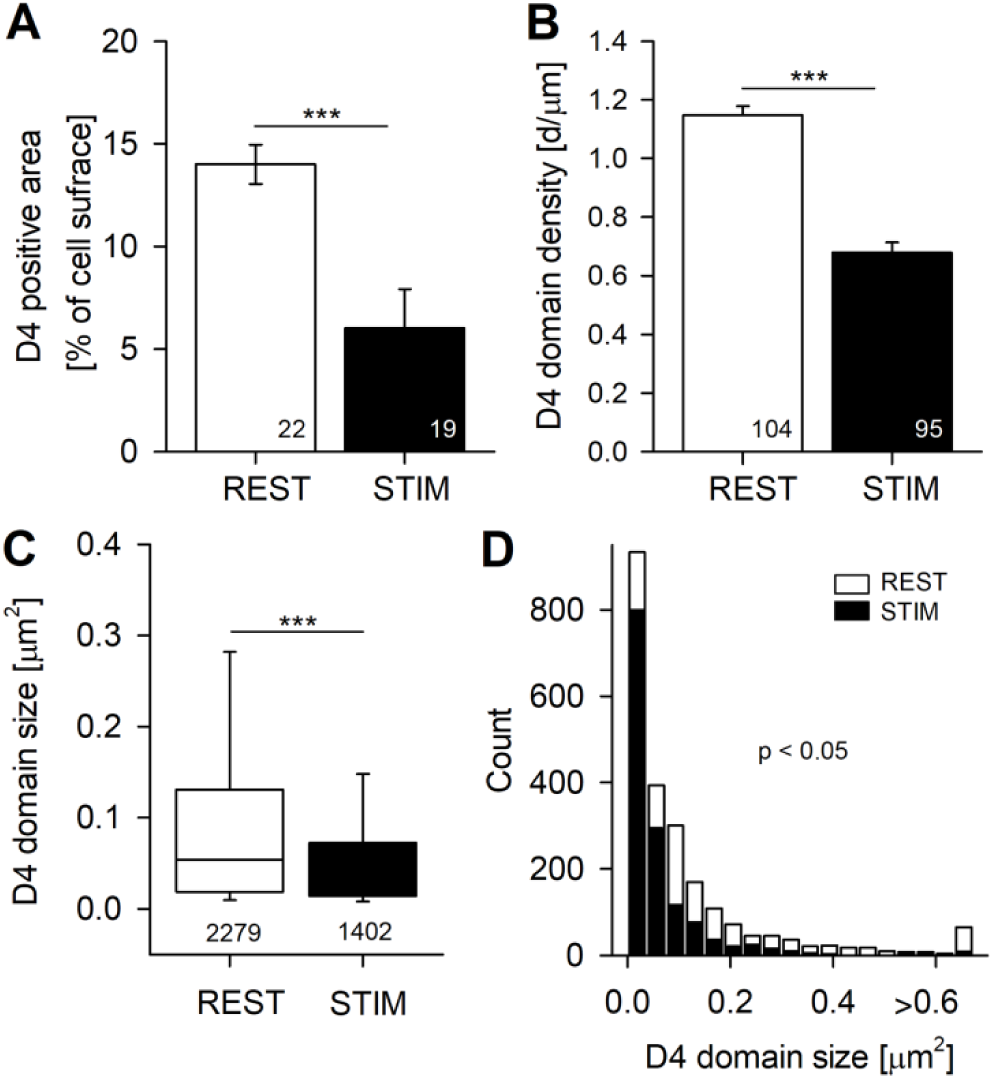
Stimulation of Regulated Exocytosis Decreases the Density and Area of Cholesterol-Rich Domains in the Outer Plasma Membrane Leaflet, Labeled by mCherry-D4-PFO (D4) in Lactotrophs, Related to Figure 2. We repeated the experiment shown in Figure 2E and adjusted the acquisition parameters for faint signals. (A) Relative D4-positive plasmalemmal (PM) area decreased significantly from 14.0% ± 0.9% in controls (REST, n = 22) to 6.0% ± 1.9% in exocytosis-stimulated cells (STIM, n = 19, p < 0.001). (B) The density of D4-positive domains, expressed as the number of D4-positive structures (d) per PM length (in µm), also significantly decreased from 1.15 ± 0.03 d/µm in controls (REST, n = 104 optical slices, 12 cells) to 0.68 ± 0.04 d/µm in exocytosis-stimulated cells (STIM, n = 95 optical slices, 13 cells, p < 0.001). (C) The size of the individual D4 domains (insets in Figures 2C and 2D) was significantly smaller and decreased from 0.054 µm^2^ (interquartile range [IQR], 0.019–0.131, n = 2279 domains, 12 cells) in controls (REST) to 0.032 µm^2^ (IQR, 0.014–0.072, n = 1402 domains, 13 cells) in exocytosis-stimulated cells (STIM, p < 0.001). (D) Frequency distribution of D4 domain size in vehicle-treated (REST) and stimulated (STIM; 18 bins; bin width, 0.037 μm^2^) cells. Note the significant difference in distributions (p < 0.05, Kolmogorov-Smirnov test). The numbers in the columns indicate the number of cells analyzed in (A), the number of optical slices analyzed in (B), and the number of D4 domains analyzed in (C). Data are presented as means ± SEM in (A) and (B), and as median with IQR in (C), ***p < 0.001. ANOVA with Holm-Sidak post-hoc test in (A), student’s t-test in (B), and Mann-Whitney rank sum test in (C).

**Figure S4.**
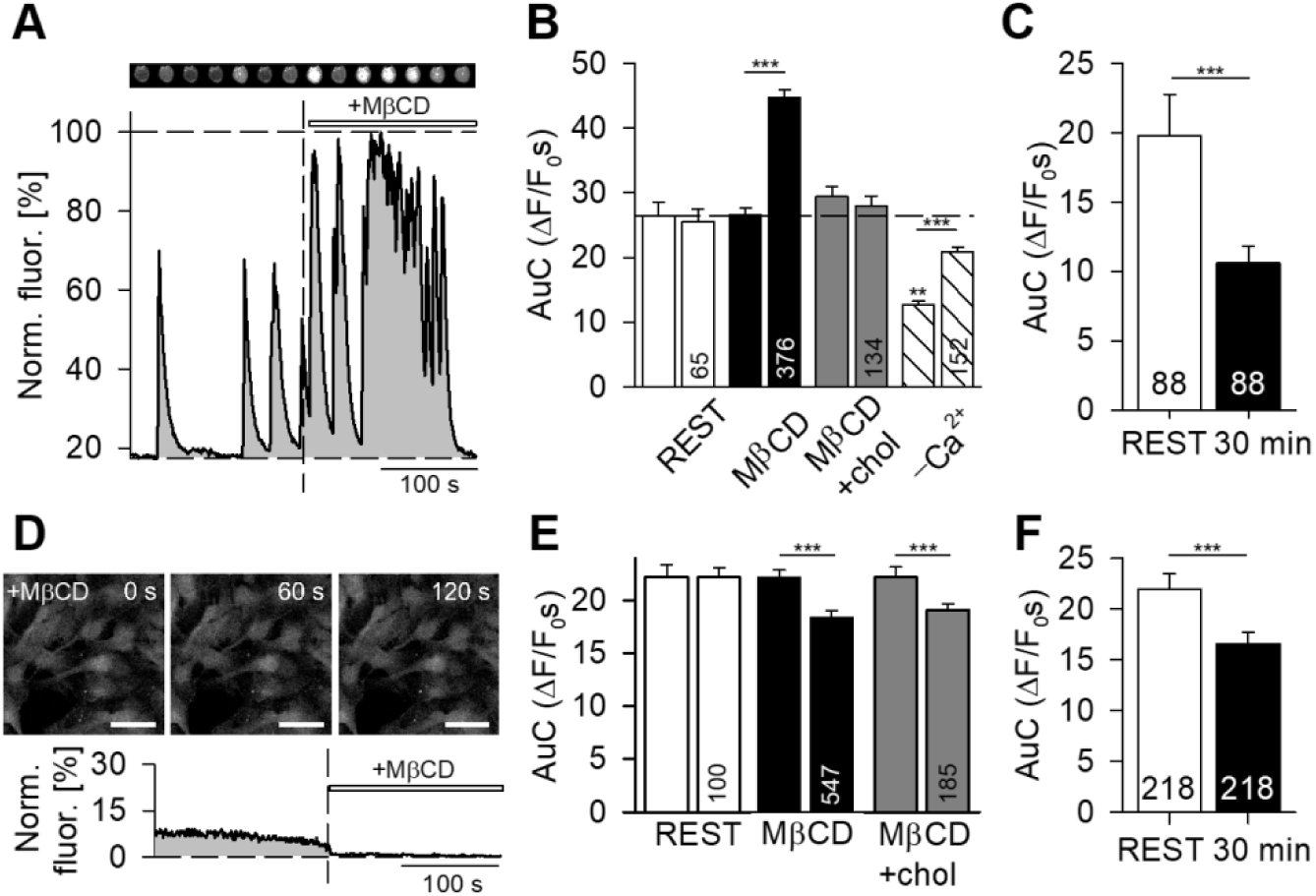
Cholesterol Extraction Increases Cytosolic Free Calcium Levels ([Ca^2+^]_i_) in Lactotrophs and Decreases [Ca^2+^]_i_ in Astrocytes, Related to Figure 4. (A) Representative confocal images of a Fluo-4-loaded ([Ca^2+^]_i_ marker) lactotroph (top sequence) and time-dependent fluctuations in [Ca^2+^]_i_ (Norm. fluor. [%]) 3 min before (before the vertical dashed line) and after MβCD treatment (10 mM, 3 min, after the vertical dashed line). (B) The area under the curve (AuC, see the gray area in A; ΔF/F_0_s) 3 min before and 3 min after treating lactotrophs with vehicle (REST), MβCD, cholesterol-replenishing solution (MβCD + chol) and Ca^2+^-free vehicle (-Ca^2+^). Dashed line, resting baseline [Ca^2+^]_i_. (C) The AuC (ΔF/F_0_s) in resting (REST) and MβCD-terated (30 min) lactotrophs. (D) Representative images of [Ca^2+^]_i_ levels in astrocytes at the time of MβCD application (+MβCD, 0 s) as well as 60 s and 120 s in the presence of MβCD; and (below) the time-dependent fluctuations in [Ca^2+^]_i_ (Norm. fluor. [%]) during resting conditions (3 min) and the same cell after acute treatment with MβCD (+MβCD, 3 min). Note the reduction in [Ca^2+^]_i_. (E) The AuC (ΔF/F_0_s) 3 min before and after treatment of astrocytes with vehicle (REST) and MβCD and cholesterol-replenishing solution (MβCD + chol). (F) The AuC (ΔF/F_0_s) in resting (REST) astrocytes and MβCD-terated (30 min) astrocytes. The numbers in the bars indicate the number of cells analyzed. Data are presented as means ± SEM, ***p < 0.001; Student’s t-test. Scale bar, 20 μm in (D).

**Figure S5.**
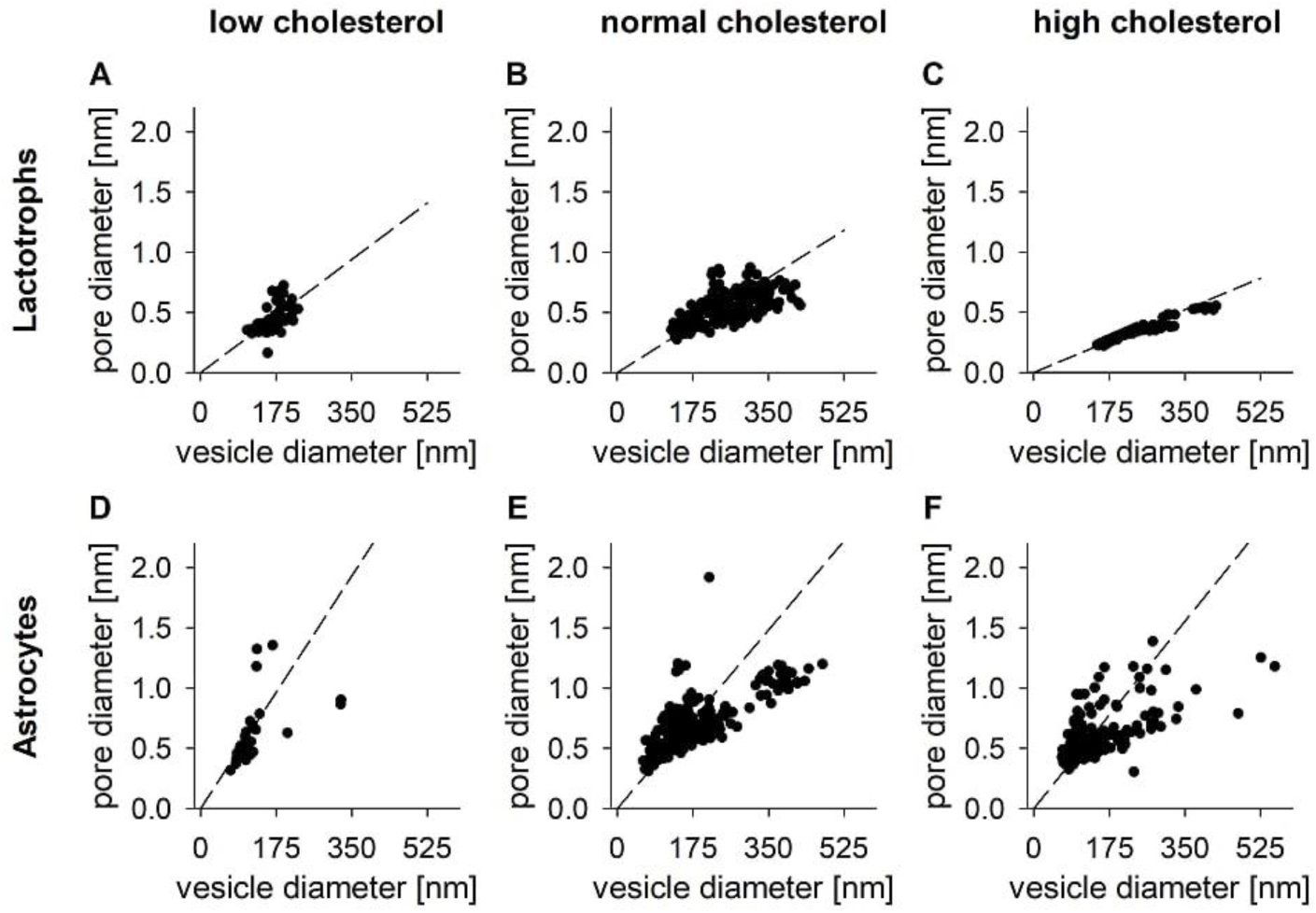
Theoretical Model Fitting Experiment, Related to Figure 5 and Figure 6. The relationship between the measured values of the fusion pore diameter (estimated from the fusion pore conductance (Gp); pore diameter [nm]) and the vesicle diameter [nm], estimated from the vesicle membrane capacitance (Cv) at different cholesterol densities (low, normal and high cholesterol). The dashed lines are the best fits using eq. 9 (Supplemental Information) with values of β/*E*_0_ denoting line slopes, which were significantly higher at low cholesterol than at normal and high cholesterol (p < 0.05, ANCOVA, Bonferroni corrected), both in lactotrophs and astrocytes.

**Figure S6.**
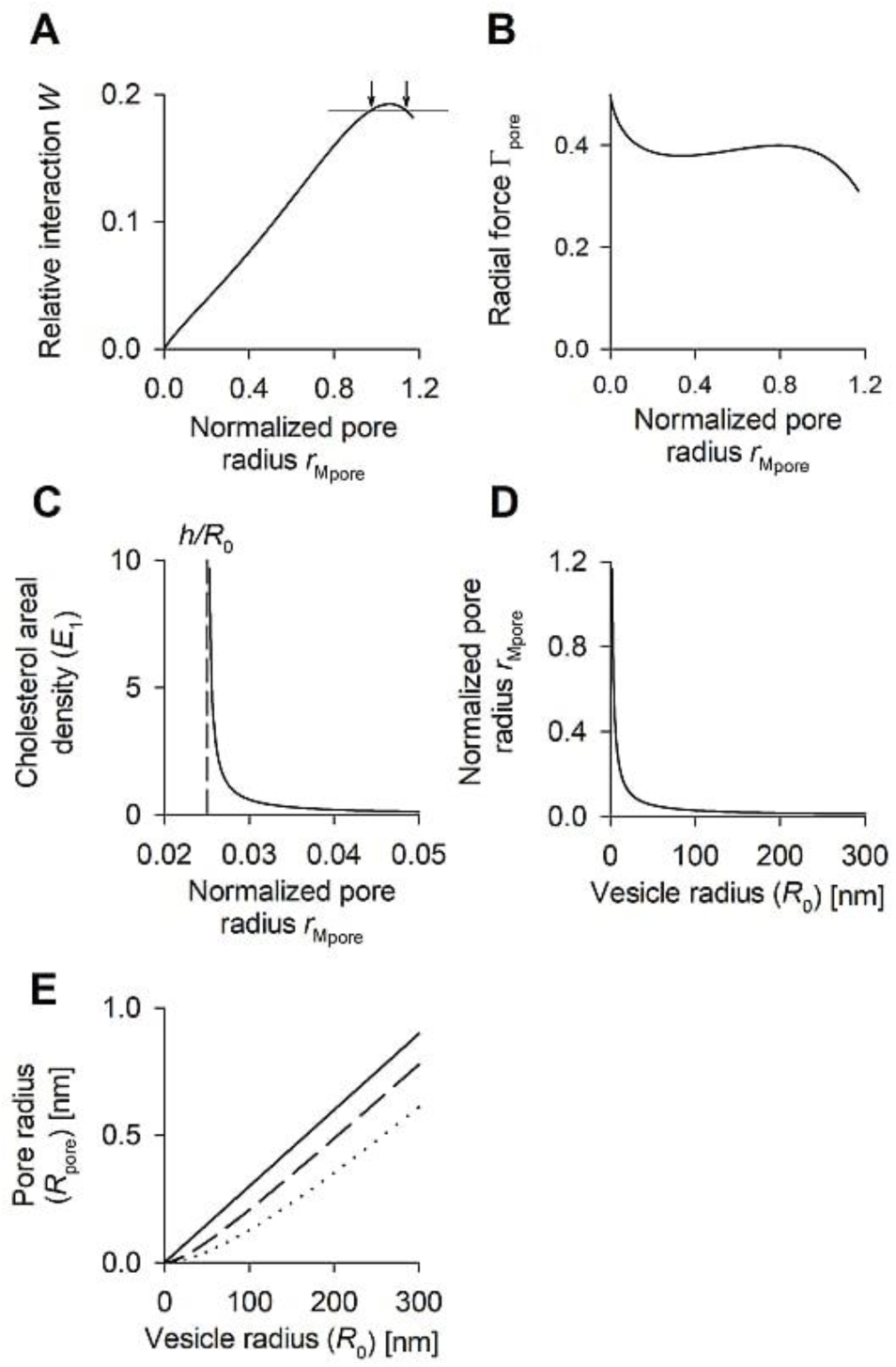
A description of the Fusion Pore Radius as a Function of the Secretory Vesicle Radius and Membrane Cholesterol Density (Concentration), Related to Figure 5. (A) The dimensionless, relative interaction constant (*W*) as a function of the normalized pore radius of the membrane midplane at the fusion pore (r_Mpore_). The arrows denote two solutions for the fusion pore radius at a given value of *W* (gray line). (B) The dimensionless transfer shear force in the radial direction (radial force, Γ_pore_) as a function of the normalized pore radius of the membrane midplane at the fusion pore (*r*_Mpore_). (C) Dependence of the cholesterol areal density in the invagination membrane (*E*_1_), relative to the corresponding plasmalemmal density, as a function of the normalized pore radius of the membrane midplane at the fusion pore (r_Mpore_) at β = 3 × 10^−3^ × *E*_0_ and h/*R*_0 =_ 0.025. (D) Dependence of the normalized pore radius of the membrane midplane at the fusion pore (*r*_Mpore_) as a function of the vesicle radius (*R*_0_) at α = 1 nm^−1^/*E*_0_, and β = 3 × 10^−3^ × *E*_0_. (E) Dependence of the fusion pore radius (*R*_pore_) as a function of the vesicle radius (*R*_0_) for the parameter α = 1 nm^−1^/*E*_0_ (full line), 0.01 nm^−1^/E_0_ (dashed line), and 0.0033 nm^−1^/E_0_ (dotted line) at β = 3 × 10^−3^ × *E*_0_. Lines in (A), (B), (D), and (E) end where the fusion pores are completely open. See Supplemental Information for the details of the model.

## Methods

## LEAD CONTACT AND MATERIALS AVAILABILITY

Further information and requests for resources and reagents should be directed to and will be fulfilled by the Lead Contact, Robert Zorec.

## EXPERIMENTAL MODEL AND SUBJECT DETAILS

### Materials and Solutions

Methyl-β-cyclodextrin (MβCD) and cholesterol were from Sigma-Aldrich. The cholesterol assay kit (Amplex Red) and Vybrant DiD cell-labeling solution were from Invitrogen. All other chemicals were of the highest purity available. The extracellular (bath) solution (ECS) for lactotrophs contained 130 mM NaCl, 5 mM KCl, 8 mM CaCl_2_, 10 mM D-glucose, 10 mM HEPES (N-2-hydroxyethylpiperazine-N′-2-ethanesulfonic acid) at pH 7.2 (with NaOH). The ECS for astrocytes contained 130 mM NaCl, 5 mM KCl, 2 mM CaCl_2_, 1 mM MgCl_2_, 10 mM D-glucose, 10 mM HEPES at pH 7.2 (with NaOH). In experiments with Ca^2+^-deficient extracellular solution, CaCl_2_ was replaced with NaCl. The osmolarity of the ECS was ∼300 mOsm.

### Animals

The experiments were performed on primary lactotroph cultures isolated from 30- to 60-day-old Wistar rats and primary astrocyte cultures isolated from 2- to 3-day-old Wistar rats. The care of the experimental animals was in accordance with the International Guiding Principles for Biomedical Research Involving Animals developed by the Council for International Organizations of Medical Sciences and Directive on Conditions for Issue of License for Animal Experiments for Scientific Research Purposes (Official Gazette of the RS, nos. 40/85 and 22/87). The research conducted was approved by The Administration of the Republic of Slovenia for Food Safety, Veterinary and Plant Protection (Republic of Slovenia, Ministry of Agriculture, Forestry and Food, Dunajska cesta 22, 1000 Ljubljana), document no. U34401-47/2014/7.

### Cell Cultures

Lactotroph primary cultures were prepared as described (Rituper et al., 2013b). After isolation, lactotrophs were plated onto poly-L-lysine (1%, w/v)-coated coverslips and kept in high-glucose Dulbecco’s modified Eagle’s medium (Invitrogen) supplemented with 10% newborn calf serum and L-glutamine at 37°C with 95% humidity and 5% CO_2_. Primary cortical astrocyte cultures were prepared from the cerebral cortices of neonatal rats as described (Schwartz and Wilson, 1992). Cells were grown in Dulbecco’s modified Eagle’s medium supplemented with 10% fetal bovine serum, 1 mM sodium pyruvate, 2 mM l-glutamine, and 5 U/mL penicillin / 5 μg/mL streptomycin at 37°C with 95 % humidity and 5% CO_2_. When cells reached confluence, they were shaken overnight, trypsinized, and plated into 2-mL cell culture tubes. After reaching confluence again, they were subcultured onto 22-mm poly-L-lysine-coated (0.1 %, w/v) coverslips. Experiments were performed on lactotrophs and astrocytes within 3 days after plating.

Fibroblasts were prepared from *Npc1* null mice. Briefly, BALB/cNctr-Npc1m1N/J, (*Npc1*^*-/*-^) and control (*Npc1*^*+/+*^) mice were generated from heterozygous breeding. Genotyping was performed according to published methods (Loftus et al., 1997). All mice were maintained under a standard 12h light/12h dark cycle with water and food available ad libitum. All procedures were performed according to the UK Animals (Scientific Procedures) Act 1986 under a project license (PPL No. I25BD461E) from the UK Home Office. Fibroblasts were prepared from *Npc1*^*-/-*^ and *Npc1*^*+/+*^ mice according to published methods (Khan and Gasser, 2016).

## METHOD DETAILS

### Fluorescent (TopFluor) Cholesterol Loading

Fluorescent cholesterol (TopFluor cholesterol)-MβCD inclusion complexes were prepared as described (Rituper et al., 2013a). Briefly, 10 mg of TopFluor cholesterol (23-(dipyrromethene boron difluoride)-24-norcholesterol) (Avanti Polar Lipids) was added to the 10 mM MβCD solution (in phosphate buffered saline [PBS], molar ratio of MβCD/cholesterol 8:1) and sonicated in an ice bath (4°C) for 30 min protected from light. The solution was filtered through 0.2-µm filters, aliquoted, and stored at −20°C until further use. To load fluorescent cholesterol into prolactin (PRL) secretory vesicles, lactotroph-loaded coverslips were washed with PBS and incubated with 1 mL of loading solution, diluted to a TopFluor cholesterol concentration of 50 μg/mL at 37°C and 5% CO_2_ for 60 min. After thorough washing (in PBS), loading solution was replaced with standard lactotroph culture medium for 8−10 h. High background fluorescence could be diminished by a brief (2 min) incubation of cells with 10 mM MβCD solution.

### Immunocytochemistry

PRL granules attached to the cell surface were immunolabeled by first blocking nonspecific immunoreactive sites with 3% bovine serum albumin (BSA/PBS) and then tagging the PRL antigens with primary mouse monoclonal anti-PRL antibodies (1:300; Thermo Scientific) for 10 min at room temperature (RT). To immunolabel PRL secretory vesicles, cells were fixed and permeabilized with 2% formaldehyde (10 min, RT) and incubated in 3% BSA and 10% goat serum in PBS (1 h, 37°C). PRL vesicles were labeled with primary mouse monoclonal anti-PRL antibodies (1:300, Thermo Scientific) for 2 h at 37°C. Primary anti-PRL antibodies were localized using secondary anti-mouse antibodies conjugated to specified Alexa fluorophores (1:600, 45 min, 37°C). Glutamatergic and ATP-containing vesicles were immunolabeled by fixing astrocytes with 4% formaldehyde (15 min, RT). Astrocytes were then incubated in 3% BSA and 10% goat serum in PBS (1 h, 37°C). Next, primary mouse monoclonal anti-VGLUT1 antibodies (1:800; SySy) were added for 2 h at 37°C and then primary rabbit polyclonal anti-VNUT antibodies (1:200, from Prof. Yoshinori Moriyama) for 2 h at 37°C, followed by the addition of both secondary antibodies conjugated to specified Alexa fluorophores (1:600, 45 min, 37°C). SlowFade Gold antifade reagent was used to mount coverslips to glass slides (Invitrogen).

### EGFP-D4-PFO- and mCherry-D4-PFO Expression and Purification

The perfringolysin O (PFO) D4 domain with N-terminal EGFP (EGFP-D4-PFO) or mCherry (mCherry-D4-PFO) fusion was expressed and purified as described previously (Lasic et al., 2019). Briefly, an EGFP- or mCherry-tagged D4 DNA fragment coding for the amino acids from S386 to N500 was introduced into the pET8c vector using XhoI and MluI sites. The resulting expression plasmid encoded for N-terminal 6xHis-tag followed by the recognition site for TEV protease and the EGFP/mCherry coding region separated from the D4 PFO domain with 10 amino acids long GS linker. The integrity of each plasmid was verified by DNA sequencing of its entire open reading frame. For recombinant protein production, a freshly transformed *Escherichia coli* BL21(DE3) strain was used, which was grown at 37°C in LB medium supplemented with 100 µg/mL ampicillin. Overexpression of the fluorescently tagged D4 domain was induced with 0.5 mM isopropyl β-D-1-thiogalactopyranoside (IPTG) at OD_600_ from 0.5 to 0.7 for 16−18 h at 20°C. The cells were harvested by centrifugation, resuspended in 10 mL/g wet mass of lysis buffer (50 mM NaH_2_PO4, 300 mM NaCl, 10 mM imidazol [pH 8.0]) and lysed by sonication. Recombinant proteins were purified by nickel chromatography (Ni-NTA Superflow; Qiagen) using the same procedures as described previously (Lasic et al., 2019). Purified protein was dialyzed against Tris-HCl buffer (50 mM Tris-HCl, 200 mM NaCl, 5% (v/v) glycerol [pH 7.4]) using Slide-a-lyzer with a molecular weight cutoff of 10 kDa (Thermo Scientific). Purification was confirmed by SDS-PAGE and Coomassie staining.

### D4-PFO Membrane and Organelle Labeling, and D4-PFO Expression in Mammalian Cells

To label the exoplasmic leaflet of lactotroph PM, cells were first rinsed with PBS followed by incubation with 1 µM mCherry-D4-PFO or EGFP-D4-PFO in 3% BSA for 30 min at RT. After washing, cells were fixed with 2% formaldehyde. In some experiments, cells were further immunolabeled as described in the Results (Figure 3). In addition to the labeling, the mCherry-D4-PFO-encoding plasmid was also expressed in lactotrophs. For this purpose, gene fragment coding for mCherry-D4 fusion from the bacterial expression plasmid described above was amplified with the following pair of primers: 5′-CGT ACA AGC TTA TGC ATC ACC ATC ACC ATC AC-3′ (sense) and 5′-GCA TGT GGA TCC AGT ACG CGT TTA ATT GTA AGT AAT ACT-3′ (antisense). The PCR product was cloned into the pcDNA3 vector using HindIII and BamHI restriction sites. The sequence of the constructed plasmid was confirmed by DNA sequencing. Electroporation was used to transfect plasmids into lactotrophs using a Basic Nucleofector Kit for Primary Mammalian Epithelial Cells (Lonza, program W-001) (Figure S1).

To examine the distribution of cholesterol-rich membranous organelles in fibroblasts (wild-type and NPC1^−/−^ lacking the vesicle cholesterol transporter Niemann-Pick Type C1), cells were washed (3 min) with PBS, fixed in formaldehyde (2% in PBS) for 10 min, washed three times (3 min) with PBS, exposed to D4-PFO-mCherry (0.25 µM) for 30 min, and washed three times with PBS (3 min); all at RT. Finally, coverslips were mounted onto glass slides using SlowFade Gold antifade mountant (Thermo Scientific).

#### Plasma Membrane Labeling with Vybrant DiD Solution

In some experiments, plasma membrane was labeled with 25 µM Vybrant DiD cell-labeling solution (Invitrogen) for 10 min at RT. Cells were then fixed in 2% formaldehyde and mounted on glass slides using SlowFade Gold antifade reagent (Invitrogen).

## QUANTIFICATION AND STATISTICAL ANALYSIS

### Electrophysiological Measurements

Cell-attached capacitance measurements were performed with a dual-phase lock-in patch-clamp amplifier (SWAM IIC; Celica) as described (Rituper et al., 2013b). All experiments were performed at RT. Individual recordings lasted 1,000 s. We used fire-polished pipettes, heavily coated with Sylgard and with a tip resistance of 2–5 MΩ. The bath and pipettes contained ECS. A sine wave (111 mV rms and 1591 Hz for lactotrophs and 111 mV rms and 6364 Hz for astrocytes) was applied to the pipette, and the pipette potential was held at 0 mV. During the experiments, the phase angle was adjusted to nullify the changes in the real (Re) trace in response to the manually generated 10 fF calibration pulses. Vesicle capacitance (C_v_) and fusion pore conductance (G_p_) were calculated from the imaginary (Im) and real (Re) parts of the admittance traces as reported previously (Lollike and Lindau, 1999): C_v_ = [(Re^2^ + Im^2^)/Im]/ω, where ω is the angular frequency (ω = 2πf, f is the sine wave frequency) and G_p_ = (Re^2^ + Im^2^)/Re. The fusion pore radius was estimated using the equation G_p_ = (π*R*_pore_^2^)/(ρλ), where *R*_pore_ represents the fusion pore radius, ρ is the estimated resistivity of saline (100 Ω cm), and λ the estimated length of a gap junction channel (15 nm) (Spruce et al., 1990). The vesicle diameter was calculated using a specific membrane capacitance (C_sm_) of 8 fF/µm^2^ (lactotrophs) or 10 fF/µm^2^ (astrocytes) as described (Rituper et al., 2013b). We used a custom-written MATLAB (Math Works) subroutine (CellAn; Celica) to analyze fusion events.

### Total Cell Cholesterol Measurements

The total cell cholesterol (with cholesterol esters) was determined using the Amplex Red Cholesterol Assay Kit (Invitrogen). Cells were isolated as described above, however, no serum or serum substitutes were used. Samples were treated with 10 mM MβCD for 30 min at 37°C in a 5% CO_2_ atmosphere. For cholesterol replenishment, cells were washed and resuspended in 10 mM cholesterol loading solution (MβCD/cholesterol molar ratio of 8:1), prepared as described previously (Churchward et al., 2005b) for an additional 30 min. Subsequently, cells were lysed with CelLytic (Sigma-Aldrich), mixed with Amplex Red reagent, and incubated for 60 min at 37°C in a 5% CO_2_ atmosphere. The resulting fluorescence was measured using an EnSpire 2300 Multimode Plate Reader (PerkinElmer) with excitation at 560 nm and fluorescence emission detected at 590 nm.

### Prolactin Release Measurements

To measure PRL release, lactotrophs were densely plated on 22-mm coverslips and treated with vehicle or 10 mM MβCD for 30 min. Then, the bath solution was collected, and the rat PRL enzyme immunoassay (ELISA) was performed according to the manufacturer’s instructions (A05101, SPI bio; Bertin Pharma).

### ANP.emd Release Measurements

Astrocytes were transfected with the plasmid encoding atrial natriuretic peptide tagged with emerald green fluorescent protein (ANP.emd; a gift from Dr. Ed Levitan, University of Pittsburgh, Pittsburgh, PA, USA) using a Rat Astrocyte Nucleofector Kit (Lonza). Transfected cells were resuspended in culture medium and subcultured onto poly-l-lysine-coated coverslips or 24-well plates (Nunc, Thermo Scientific) and maintained at 37°C in an atmosphere of 5% CO_2_ for 48–72 h. The release of ANP.emd was estimated by measuring the fluorescence intensity of the bath solution at specified time points (EnSpire microplate reader; PerkinElmer) at the emission wavelength of 508 nm. Fluorescence was excited at 488 nm. Background fluorescence was subtracted, and the measured values were averaged and expressed as a relative increase in fluorescence, normalized to controls. A one-sample t-test was used for statistical comparison.

### Calcium Measurements

Cells were loaded with 2 µM Fluo-4 AM (Molecular Probes) for 30 min in ECS. After rinsing in ECS or Ca^2+^-free ECS for 15 min, the cells were placed into the recording chamber and imaged with a confocal laser scanning microscope (Zeiss LSM 780). Measurements were performed with a Zeiss Plan-Apochromat oil immersion objective (40×, NA 1.3). An Ar-ion laser was used to excite Fluo-4 in combination with a 505-nm long-pass emission filter. Time-lapse images were acquired at a sampling rate of 1 Hz. First, baseline fluorescence intensities were recorded for 3 min (F_0_). Then, cells were treated with either vehicle, 10 mM MβCD, or 10 mM cholesterol loading solution, and fluorescent images were acquired for an additional 3 min. To assess the effect of prolonged exposure to MβCD, fluorescence images were acquired for the initial 3 min and then for an additional 3 min after 30 min of incubation with MβCD. Fluo-4 fluorescence, which reports changes in [Ca^2+^]_i_, was measured in a region of interest covering the entire cell image. Resulting time-dependent fluorescence intensity traces were first normalized to the maximal fluorescence intensity in a given trace. The fluorescence intensity change was defined as ΔF/F_0_ = (F(t) − F_0_)/F_0_, expressed as a percentage. F_0_ is the average non-stimulated fluorescence intensity of ten frames with the lowest relative intensity of fluorescence, and F(t) is the relative fluorescence intensity at a given time (Figure S4).

### Lipid Extraction and HPTLC Analysis

Cells (astrocytes) were lysed with CelLytic M (Sigma-Aldrich) and lipids were extracted as described (Churchward et al., 2005b). Lysed cells were suspended in PBS, and CH_3_OH and CHCl_3_ were added sequentially at a ratio of 0.8:2:1 (PBS/CH_3_OH/CHCl_3_, v/v/v), followed by vortexing and sonication for 30 s. An aqueous solution of 1 M NaCl and 0.1 M HCl was added, followed by the addition of CHCl_3_ to reach a final ratio of 1.8:2:2 (PBS/CH_3_OH/CHCl_3_, v/v/v). The sample was vortexed vigorously after each addition. The lower organic phase was dried under N_2_ and dissolved in CHCl_3_/CH_3_OH (2:1) for automated high-performance thin-layer chromatography (HPTLC) using the CAMAG AMD 2 system. Silica gel 60 plates (Merck) were pre-washed with CH_3_OH/C_4_H_8_O_2_ (6:4 v/v), and heat activated at 110°C for 30 min before analysis. Neutral lipids and phospholipids were resolved as previously described (Churchward et al., 2008a), detected using copper sulfate charring, and imaged using the LAS 4000 Biomolecular Imager (GE Healthcare). Images were analyzed using Multi Gauge Software (FUJIFILM Corporation) or ImageJ (NIH). Fluorescence of control samples was defined as 100%, and changes in lipid concentrations after treatment with MβCD are reported relative to corresponding controls.

### Microscopy and Image Analysis

Except where stated otherwise, structured illumination microscopy (Zeiss Elyra PS1) was used to image fluorescently labeled cells. Images were acquired with a Zeiss Plan-neofluar oil immersion objective (63×, NA 1.4). Cells were illuminated with laser lines of 488 nm and/or 561 nm, and the emitted fluorescence was collected through band-pass emission filters (495–560 nm and 570–650 nm, respectively). Unless specified otherwise, 200-nm-thick Z stacks were acquired with an EMCCD camera (16-bit, Andor iXon 885; Andor Technology) with variable exposure and analyzed with ImageJ.

#### Analysis of D4-PFO-Labeled Plasma Membrane Domains

D4-PFO labeling and analysis protocols are depicted schematically in Figure S1. Briefly, after labeling with D4-PFO (and PRL in some experiments), single (D4-PFO) or dual channel (D4-PFO and PRL) fluorescent images were acquired and a 200 nm interval Z stack image was constructed for every cell. Then a 20-pixel-wide band covering the plasma membrane region was selected in every image within the stack (5–15 optical slices per cell), straightened, and thresholded automatically using the methods for D4-PFO (Kapur et al., 1985) and PRL labeling (Tsai, 1995). Thresholded images were further analyzed as described in Figure 3.

#### Surface-Attached PRL Cargo Granules Count

Structured illumination microscopy images within the Z stack of lactotrophs (labeled for surface PRL granules; see Table S2) were recorded at 200 nm intervals, and the resulting Z stacks were used to reconstruct imaged cells in 3D. The 3D Objects Counter plugin (Bolte and Cordelieres, 2006) for ImageJ was used to calculate the number of PRL surface granules. Results were normalized to the cell volume, which was calculated as described (Rituper et al., 2013a).

#### Estimation of Secretory Vesicle Quantity

Confocal laser scanning microscopy (Zeiss LSM 780) was used to image immunolabeled vesicles in fixed permeabilized lactotrophs and astrocytes. Images were acquired with a Zeiss Plan-Apochromat oil immersion objective (63×, NA 1.4). For excitation of Alexa Fluor 488, an Ar-ion laser was used in combination with a band-pass emission filter (505–530 nm), and for excitation of Alexa Fluor 543, a He-Ne laser was used in combination with a long-pass emission filter (cutoff below 560 nm). Imaged cells were Z sectioned at 400-nm-thick intervals. Fluorescence images were thresholded automatically using an iterative selection method (Ridler and Calvard, 1978) for lactotrophs and another method (Kapur et al., 1985) for astrocytes. The percentage cell area (in an optical slice) positive for immunolabeled vesicles was calculated by dividing above-threshold pixels with all the pixels in a region of interest covering the entire cell (cell borders were determined with the help of transmission images).

#### Analysis of D4-PFO-Labeled Membranous Organelles in Fibroblasts

To estimate the vesicle counts and vesicle intensity, confocal images were analyzed with ImageJ. The minimum fluorescent spot taken to identify an individual vesicle in a confocal image was three adjacent pixels (0.132 × 0.132 μm), and the minimum surface area covered by a spot was 0.052 µm^2^. Considering that the minimum D4-PFO-positive vesicle consisted of three adjacent pixels, a broad span of vesicles with different sizes and intensity was covered by this analysis.

### Statistical Tests

Experimental data were compared with either the Student’s t test, one-way ANOVA with Holm-Sidak’s post-hoc test or ANOVA on ranks with Dunn’s post-hoc test, as indicated. Depending on the distribution, the data are presented as means ± SEM or medians with interquartile range. Differences were considered significant at *p < 0.05, **p < 0.01, and ***p < 0.001. Statistical analyses were carried out using SigmaPlot (Systat Sofware). Data in Figure S5 were fitted using standard model linear regression, and the slopes were statistically compared with the analysis of covariance (ANCOVA) method.

## DATA AND CODE AVAILABILITY

The datasets supporting the current study are available from the corresponding author on request, as these have not been deposited in a public repository because of potential patent application.

## SUPPLEMENTAL INFORMATION

Supplemental Information can be found in the attached file.

## SUPPORTING CITATIONS

The following references appear in the Supplemental Information: (Deuling and Helfrich, 1976; Helfrich, 1973; Heuser, 1989; Julicher and Lipowsky, 1996; Julicher and Seifert, 1994; Kuchel and Ralston, 1988; Seifert et al., 1991; Svetina and Zeks, 1989).

## REFERENCES

Akerman, S.N., Zorec, R., Cheek, T.R., Moreton, R.B., Berridge, M.J., and Mason, W.T. (1991). Fura-2 imaging of thyrotropin-releasing hormone and dopamine effects on calcium homeostasis of bovine lactotrophs. Endocrinology 129, 475–488.

Almers, W., and Tse, F.W. (1990). Transmitter release from synapses: does a preassembled fusion pore initiate exocytosis? Neuron 4, 813–818.

Alvarez de Toledo, G., Fernandez-Chacon, R., and Fernandez, J.M. (1993). Release of secretory products during transient vesicle fusion. Nature 363, 554–558.

Baumgart, T., Hess, S.T., and Webb, W.W. (2003). Imaging coexisting fluid domains in biomembrane models coupling curvature and line tension. Nature 425, 821–824.

Berman, R.M., Cappiello, A., Anand, A., Oren, D.A., Heninger, G.R., Charney, D.S., and Krystal, J.H. (2000). Antidepressant effects of ketamine in depressed patients. Biol Psychiatry 47, 351–354.

Bogan, J.S., Xu, Y., and Hao, M. (2012). Cholesterol accumulation increases insulin granule size and impairs membrane trafficking. Traffic 13, 1466–1480.

Bolte, S., and Cordelieres, F.P. (2006). A guided tour into subcellular colocalization analysis in light microscopy. Journal of Microscopy-Oxford 224, 213–232.

Brose, N., Brunger, A., Cafiso, D., Chapman, E.R., Diao, J., Hughson, F.M., Jackson, M.B., Jahn, R., Lindau, M., Ma, C., et al. (2019). Synaptic vesicle fusion: today and beyond. Nat Struct Mol Biol 26, 663–668.

Chang, C.W., Chiang, C.W., and Jackson, M.B. (2017). Fusion pores and their control of neurotransmitter and hormone release. J Gen Physiol 149, 301–322.

Chen, Z., and Rand, R.P. (1997). The influence of cholesterol on phospholipid membrane curvature and bending elasticity. Biophys J 73, 267–276.

Chow, R.H., von Ruden, L., and Neher, E. (1992). Delay in vesicle fusion revealed by electrochemical monitoring of single secretory events in adrenal chromaffin cells. Nature 356, 60–63.

Churchward, M.A., Brandman, D.M., Rogasevskaia, T., and Coorssen, J.R. (2008a). Copper (II) sulfate charring for high sensitivity on-plate fluorescent detection of lipids and sterols: quantitative analyses of the composition of functional secretory vesicles. J Chem Biol 1, 79–87.

Churchward, M.A., and Coorssen, J.R. (2009). Cholesterol, regulated exocytosis and the physiological fusion machine. Biochem J 423, 1–14.

Churchward, M.A., Rogasevskaia, T., Brandman, D.M., Khosravani, H., Nava, P., Atkinson, J.K., and Coorssen, J.R. (2008b). Specific lipids supply critical negative spontaneous curvature--an essential component of native Ca2+-triggered membrane fusion. Biophys J 94, 3976–3986.

Churchward, M.A., Rogasevskaia, T., Hofgen, J., Bau, J., and Coorssen, J.R. (2005a). Cholesterol facilitates the native mechanism of Ca2+-triggered membrane fusion. J Cell Sci 118, 4833–4848.

Churchward, M.A., Rogasevskaia, T., Hofgen, J., Bau, J., and Coorssen, J.R. (2005b). Cholesterol facilitates the native mechanism of Ca2+-triggered membrane fusion. Journal of Cell Science 118, 4833–4848.

Cochilla, A.J., Angleson, J.K., and Betz, W.J. (1999). Monitoring secretory membrane with FM1-43 fluorescence. Annu Rev Neurosci 22, 1–10.

Cookson, E.A., Conte, I.L., Dempster, J., Hannah, M.J., and Carter, T. (2013). Characterisation of Weibel-Palade body fusion by amperometry in endothelial cells reveals fusion pore dynamics and the effect of cholesterol on exocytosis. J Cell Sci 126, 5490–5499.

Deuling, H.J., and Helfrich, W. (1976). Red blood cell shapes as explained on the basis of curvature elasticity. Biophys J 16, 861–868.

Flasker, A., Jorgacevski, J., Calejo, A.I., Kreft, M., and Zorec, R. (2013). Vesicle size determines unitary exocytic properties and their sensitivity to sphingosine. Mol Cell Endocrinol 376, 136–147.

Gasman, S., and Vitale, N. (2017). Lipid remodelling in neuroendocrine secretion. Biol Cell 109, 381–390.

Grabner, C.P., and Moser, T. (2018). Individual synaptic vesicles mediate stimulated exocytosis from cochlear inner hair cells. Proc Natl Acad Sci U S A 115, 12811–12816.

Gucek, A., Jorgacevski, J., Singh, P., Geisler, C., Lisjak, M., Vardjan, N., Kreft, M., Egner, A., and Zorec, R. (2016). Dominant negative SNARE peptides stabilize the fusion pore in a narrow, release-unproductive state. Cell Mol Life Sci 73, 3719–3731.

Gundersen, V., Storm-Mathisen, J., and Bergersen, L.H. (2015). Neuroglial Transmission. Physiol Rev 95, 695–726.

He, L., Wu, X.S., Mohan, R., and Wu, L.G. (2006). Two modes of fusion pore opening revealed by cell-attached recordings at a synapse. Nature 444, 102–105.

Helfrich, W. (1973). Elastic properties of lipid bilayers: theory and possible experiments. Z Naturforsch C 28, 693–703.

Heuser, J.E. (1989). Review of electron microscopic evidence favouring vesicle exocytosis as the structural basis for quantal release during synaptic transmission. Q J Exp Physiol 74, 1051–1069.

Higgins, M.E., Davies, J.P., Chen, F.W., and Ioannou, Y.A. (1999). Niemann-Pick C1 is a late endosome-resident protein that transiently associates with lysosomes and the trans-Golgi network. Mol Genet Metab 68, 1–13.

Hoglinger, D., Burgoyne, T., Sanchez-Heras, E., Hartwig, P., Colaco, A., Newton, J., Futter, C.E., Spiegel, S., Platt, F.M., and Eden, E.R. (2019). NPC1 regulates ER contacts with endocytic organelles to mediate cholesterol egress. Nat Commun 10, 4276.

Holtta-Vuori, M., Uronen, R.L., Repakova, J., Salonen, E., Vattulainen, I., Panula, P., Li, Z., Bittman, R., and Ikonen, E. (2008). BODIPY-cholesterol: a new tool to visualize sterol trafficking in living cells and organisms. Traffic 9, 1839–1849.

Ikonen, E. (2018). Mechanisms of cellular cholesterol compartmentalization: recent insights. Curr Opin Cell Biol 53, 77–83.

Jorgacevski, J., Fosnaric, M., Vardjan, N., Stenovec, M., Potokar, M., Kreft, M., Kralj-Iglic, V., Iglic, A., and Zorec, R. (2010). Fusion pore stability of peptidergic vesicles. Mol Membr Biol 27, 65–80.

Jorgacevski, J., Potokar, M., Grilc, S., Kreft, M., Liu, W., Barclay, J.W., Buckers, J., Medda, R., Hell, S.W., Parpura, V., et al. (2011). Munc18-1 tuning of vesicle merger and fusion pore properties. J Neurosci 31, 9055–9066.

Julicher, F., and Lipowsky, R. (1996). Shape transformations of vesicles with intramembrane domains. Phys Rev E Stat Phys Plasmas Fluids Relat Interdiscip Topics 53, 2670–2683.

Julicher, F., and Seifert, U. (1994). Shape equations for axisymmetric vesicles: A clarification. Phys Rev E Stat Phys Plasmas Fluids Relat Interdiscip Topics 49, 4728–4731.

Kapur, J.N., Sahoo, P.K., and Wong, A.K.C. (1985). A New Method for Gray-Level Picture Thresholding Using the Entropy of the Histogram. Computer Vision Graphics and Image Processing 29, 273–285.

Khan, M., and Gasser, S. (2016). Generating Primary Fibroblast Cultures from Mouse Ear and Tail Tissues. J Vis Exp.

Klyachko, V.A., and Jackson, M.B. (2002). Capacitance steps and fusion pores of small and large-dense-core vesicles in nerve terminals. Nature 418, 89–92.

Kreft, M., Jorgacevski, J., Stenovec, M., and Zorec, R. (2018). Angstrom-size exocytotic fusion pore: Implications for pituitary hormone secretion. Mol Cell Endocrinol 463, 65–71.

Kreft, M., Stenovec, M., Rupnik, M., Grilc, S., Krzan, M., Potokar, M., Pangrsic, T., Haydon, P.G., and Zorec, R. (2004). Properties of Ca(2+)-dependent exocytosis in cultured astrocytes. Glia 46, 437–445.

Krzan, M., Stenovec, M., Kreft, M., Pangrsic, T., Grilc, S., Haydon, P.G., and Zorec, R. (2003). Calcium-dependent exocytosis of atrial natriuretic peptide from astrocytes. J Neurosci 23, 1580–1583.

Kuchel, P.W., and Ralston, B.G. (1988). Theory and Problems of Biochemistry, Vol 7 (New York Schaum’s Outline/McGraw-Hill).

Lasic, E., Lisjak, M., Horvat, A., Bozic, M., Sakanovic, A., Anderluh, G., Verkhratsky, A., Vardjan, N., Jorgacevski, J., Stenovec, M., et al. (2019). Astrocyte Specific Remodeling of Plasmalemmal Cholesterol Composition by Ketamine Indicates a New Mechanism of Antidepressant Action. Sci Rep 9, 10957.

Lasic, E., Rituper, B., Jorgacevski, J., Kreft, M., Stenovec, M., and Zorec, R. (2016). Subanesthetic doses of ketamine stabilize the fusion pore in a narrow flickering state in astrocytes. J Neurochem 138, 909–917.

Levitan, I., Fang, Y., Rosenhouse-Dantsker, A., and Romanenko, V. (2010). Cholesterol and ion channels. Subcell Biochem 51, 509–549.

Li, N., Lee, B., Liu, R.J., Banasr, M., Dwyer, J.M., Iwata, M., Li, X.Y., Aghajanian, G., and Duman, R.S. (2010). mTOR-dependent synapse formation underlies the rapid antidepressant effects of NMDA antagonists. Science 329, 959–964.

Liscum, L. (2000). Niemann-Pick type C mutations cause lipid traffic jam. Traffic 1, 218–225.

Loftus, S.K., Morris, J.A., Carstea, E.D., Gu, J.Z., Cummings, C., Brown, A., Ellison, J., Ohno, K., Rosenfeld, M.A., Tagle, D.A., et al. (1997). Murine model of Niemann-Pick C disease: mutation in a cholesterol homeostasis gene. Science 277, 232–235.

Lollike, K., Borregaard, N., and Lindau, M. (1995). The exocytotic fusion pore of small granules has a conductance similar to an ion channel. J Cell Biol 129, 99–104.

Lollike, K., and Lindau, M. (1999). Membrane capacitance techniques to monitor granule exocytosis in neutrophils. J Immunol Methods 232, 111–120.

Maekawa, M. (2017). Domain 4 (D4) of Perfringolysin O to Visualize Cholesterol in Cellular Membranes-The Update. Sensors (Basel) 17.

Mahadeo, M., Furber, K.L., Lam, S., Coorssen, J.R., and Prenner, E.J. (2015). Secretory vesicle cholesterol: Correlating lipid domain organization and Ca2+ triggered fusion. Biochim Biophys Acta 1848, 1165–1174.

Mason, W.T., Rawlings, S.R., Cobbett, P., Sikdar, S.K., Zorec, R., Akerman, S.N., Benham, C.D., Berridge, M.J., Cheek, T., and Moreton, R.B. (1988). Control of secretion in anterior pituitary cells--linking ion channels, messengers and exocytosis. J Exp Biol 139, 287–316.

Mukherjee, S., and Maxfield, F.R. (2004). Lipid and cholesterol trafficking in NPC. Biochim Biophys Acta 1685, 28–37.

Mukherjee, S., Soe, T.T., and Maxfield, F.R. (1999). Endocytic sorting of lipid analogues differing solely in the chemistry of their hydrophobic tails. J Cell Biol 144, 1271–1284.

Najafinobar, N., Mellander, L.J., Kurczy, M.E., Dunevall, J., Angerer, T.B., Fletcher, J.S., and Cans, A.S. (2016). Cholesterol Alters the Dynamics of Release in Protein Independent Cell Models for Exocytosis. Sci Rep 6, 33702.

Nanavati, C., Markin, V.S., Oberhauser, A.F., and Fernandez, J.M. (1992). The exocytotic fusion pore modeled as a lipidic pore. Biophys J 63, 1118–1132.

Neher, E., and Marty, A. (1982). Discrete changes of cell membrane capacitance observed under conditions of enhanced secretion in bovine adrenal chromaffin cells. Proc Natl Acad Sci U S A 79, 6712–6716.

Ohno-Iwashita, Y., Iwamoto, M., Ando, S., Mitsui, K., and Iwashita, S. (1990). A modified theta-toxin produced by limited proteolysis and methylation: a probe for the functional study of membrane cholesterol. Biochim Biophys Acta 1023, 441–448.

Rand, R.P., and Parsegian, V.A. (1986). Mimicry and mechanism in phospholipid models of membrane fusion. Annu Rev Physiol 48, 201–212.

Rawicz, W., Olbrich, K.C., McIntosh, T., Needham, D., and Evans, E. (2000). Effect of chain length and unsaturation on elasticity of lipid bilayers. Biophys J 79, 328–339.

Ridler, T.W., and Calvard, S. (1978). Picture Thresholding Using an Iterative Selection Method. Ieee Transactions on Systems Man and Cybernetics 8, 630–632.

Rituper, B., Chowdhury, H.H., Jorgacevski, J., Coorssen, J.R., Kreft, M., and Zorec, R. (2013a). Cholesterol-mediated membrane surface area dynamics in neuroendocrine cells. Biochim Biophys Acta 1831, 1228–1238.

Rituper, B., Flašker, A., Guček, A., Chowdhury, H.H., and Zorec, R. (2012). Cholesterol and regulated exocytosis: A requirement for unitary exocytotic events. Cell Calcium 52, 250–258.

Rituper, B., Gucek, A., Jorgacevski, J., Flasker, A., Kreft, M., and Zorec, R. (2013b). High-resolution membrane capacitance measurements for the study of exocytosis and endocytosis. Nat Protoc 8, 1169–1183.

Rogasevskaia, T., and Coorssen, J.R. (2006). Sphingomyelin-enriched microdomains define the efficiency of native Ca(2+)-triggered membrane fusion. J Cell Sci 119, 2688–2694.

Sanacora, G., Treccani, G., and Popoli, M. (2012). Towards a glutamate hypothesis of depression: an emerging frontier of neuropsychopharmacology for mood disorders. Neuropharmacology 62, 63–77.

Schwartz, J.P., and Wilson, D.J. (1992). Preparation and characterization of type 1 astrocytes cultured from adult rat cortex, cerebellum, and striatum. Glia 5, 75–80.

Seifert, U., Berndl, K., and Lipowsky, R. (1991). Shape transformations of vesicles: Phase diagram for spontaneous-curvature and bilayer-coupling models. Phys Rev A 44, 1182–1202.

Sharma, S., and Lindau, M. (2018). The fusion pore, 60 years after the first cartoon. FEBS Lett.

Shin, W., Arpino, G., Thiyagarajan, S., Su, R., Ge, L., McDargh, Z., Guo, X., Wei, L., Shupliakov, O., Jin, A., et al. (2020). Vesicle Shrinking and Enlargement Play Opposing Roles in the Release of Exocytotic Contents. Cell Rep 30, 421–431 e427.

Shin, W., Ge, L., Arpino, G., Villarreal, S.A., Hamid, E., Liu, H., Zhao, W.D., Wen, P.J., Chiang, H.C., and Wu, L.G. (2018). Visualization of Membrane Pore in Live Cells Reveals a Dynamic-Pore Theory Governing Fusion and Endocytosis. Cell 173, 934–945 e912.

Sikdar, S.K., Zorec, R., Brown, D., and Mason, W.T. (1989). Dual effects of G-protein activation on Ca-dependent exocytosis in bovine lactotrophs. FEBS Lett 253, 88–92.

Skocaj, M., Resnik, N., Grundner, M., Ota, K., Rojko, N., Hodnik, V., Anderluh, G., Sobota, A., Macek, P., Veranic, P., et al. (2014). Tracking cholesterol/sphingomyelin-rich membrane domains with the ostreolysin A-mCherry protein. PLoS One 9, e92783.

Spruce, A.E., Breckenridge, L.J., Lee, A.K., and Almers, W. (1990). Properties of the fusion pore that forms during exocytosis of a mast cell secretory vesicle. Neuron 4, 643–654.

Stenovec, M., Kreft, M., Grilc, S., Pangrsic, T., and Zorec, R. (2008). EAAT2 density at the astrocyte plasma membrane and Ca(2 +)-regulated exocytosis. Mol Membr Biol 25, 203–215.

Stenovec, M., Kreft, M., Poberaj, I., Betz, W.J., and Zorec, R. (2004). Slow spontaneous secretion from single large dense-core vesicles monitored in neuroendocrine cells. FASEB J 18, 1270–1272.

Stenovec, M., Lasic, E., Bozic, M., Bobnar, S.T., Stout, R.F., Jr., Grubisic, V., Parpura, V., and Zorec, R. (2016). Ketamine Inhibits ATP-Evoked Exocytotic Release of Brain-Derived Neurotrophic Factor from Vesicles in Cultured Rat Astrocytes. Mol Neurobiol 53, 6882–6896.

Stenovec, M., Trkov, S., Kreft, M., and Zorec, R. (2014). Alterations of calcium homoeostasis in cultured rat astrocytes evoked by bioactive sphingolipids. Acta Physiol (Oxf) 212, 49–61.

Stratton, B.S., Warner, J.M., Wu, Z., Nikolaus, J., Wei, G., Wagnon, E., Baddeley, D., Karatekin, E., and O’Shaughnessy, B. (2016). Cholesterol Increases the Openness of SNARE-Mediated Flickering Fusion Pores. Biophys J 110, 1538–1550.

Svetina, S., and Zeks, B. (1989). Membrane bending energy and shape determination of phospholipid vesicles and red blood cells. Eur Biophys J 17, 101–111.

Tian, A., and Baumgart, T. (2009). Sorting of lipids and proteins in membrane curvature gradients. Biophys J 96, 2676–2688.

Trkov, S., Stenovec, M., Kreft, M., Potokar, M., Parpura, V., Davletov, B., and Zorec, R. (2012). Fingolimod--a sphingosine-like molecule inhibits vesicle mobility and secretion in astrocytes. Glia 60, 1406–1416.

Tsai, W.-H. (1995). Moment-preserving thresholding: a new approach. In Document image analysis, O.G. Lawrence, and K. Rangachar, eds. (IEEE Computer Society Press), pp. 44–60.

Vardjan, N., Stenovec, M., Jorgacevski, J., Kreft, M., and Zorec, R. (2007). Subnanometer fusion pores in spontaneous exocytosis of peptidergic vesicles. J Neurosci 27, 4737–4746.

Wang, N., Kwan, C., Gong, X., de Chaves, E.P., Tse, A., and Tse, F.W. (2010). Influence of cholesterol on catecholamine release from the fusion pore of large dense core chromaffin granules. J Neurosci 30, 3904–3911.

Wasser, C.R., Ertunc, M., Liu, X., and Kavalali, E.T. (2007). Cholesterol-dependent balance between evoked and spontaneous synaptic vesicle recycling. J Physiol 579, 413–429.

Winkler, M.B.L., Kidmose, R.T., Szomek, M., Thaysen, K., Rawson, S., Muench, S.P., Wustner, D., and Pedersen, B.P. (2019). Structural Insight into Eukaryotic Sterol Transport through Niemann-Pick Type C Proteins. Cell 179, 485–497 e418.

Wustner, D., and Solanko, K. (2015). How cholesterol interacts with proteins and lipids during its intracellular transport. Biochim Biophys Acta 1848, 1908–1926.

Xu, Y., Toomre, D.K., Bogan, J.S., and Hao, M. (2017). Excess cholesterol inhibits glucose-stimulated fusion pore dynamics in insulin exocytosis. J Cell Mol Med 21, 2950–2962.

Yesylevskyy, S.O., Rivel, T., and Ramseyer, C. (2017). The influence of curvature on the properties of the plasma membrane. Insights from atomistic molecular dynamics simulations. Sci Rep 7, 16078.

Zorec, R., Sikdar, S.K., and Mason, W.T. (1991). Increased cytosolic calcium stimulates exocytosis in bovine lactotrophs. Direct evidence from changes in membrane capacitance. J Gen Physiol 97, 473–497.

