## Supplemental info on Cholesterol-mediated fusion pore regulation for "Redistribution of cholesterol from vesicle to plasmalemma controls fusion pore geometry"

### Supplemental Information

#### Table S1. Labeling Protocols with mCherry-D4-PFO (D4) and Prolactin Antibodies, Related to STAR Methods.

| Labeling protocol | A | B | C | D | E | F | G |
|---|---|---|---|---|---|---|---|
| mCherry-D4-PFO labeling | + | + | + | + | + | + | + |

| | | | | | | | |
|---|---|---|---|---|---|---|---|
| Vehicle | + | | + | | | + | |
| STIM (50 mM KCl) | | + | | + | + | | + |
| fix and permeabilize (2 % para) | + | + | + | | | | |
| PRL antibody | | | + | + | | | |
| PRL antibody + D4 | | | | | + | | |
| EGFP-D4-PFO labeling | | | | | | + | + |
| fix (2 % para) | | | | + | + | + | + |

(A) Protocol for labeling cholesterol-rich domains in the outer leaflet of the plasmalemma in resting lactotrophs (D4 → vehicle → fixation): Figures 2A–C and 3G.
(B) Protocol for labeling cholesterol-rich domains in the outer leaflet of the plasmalemma in stimulated lactotrophs (D4 → 50 mM KCl → fixation): Figures 2D and 3G.
(C) Protocol for labeling cholesterol-rich domains in the outer leaflet of the plasmalemma and PRL-containing vesicles of resting lactotrophs (D4 → vehicle → permeabilization → PRL Ab → fixation): Figure 3A.
(D) Protocol for labeling cholesterol-rich domains in the outer leaflet of the plasmalemma and surface-attached PRL vesicle cargo of stimulated lactotrophs (D4 → 50 mM KCl → PRL Ab → fixation): Figures 3B and S1.
(E) Protocol for labeling cholesterol-rich domains, including the labeling of cholesterol-rich domains incorporated into the plasmalemma due to stimulated regulated exocytosis in lactotrophs (D4 → 50 mM KCl → PRL Ab + D4 → fixation): Figure 3C.
(F) Protocol for labeling cholesterol-rich domains, where domains are first saturated with D4 and then additionally labeled with EGFP-D4 (green fluorescent peptide) after application of vehicle.
(G) Protocol for labeling cholesterol-rich domains, where domains are first saturated with D4 and then additionally labeled with EGFP-D4 after stimulation of exocytosis with a solution containing 50 mM KCl.

### Table S2. Relative Normalized Cholesterol Concentration in MβCD and MβCD + Cholesterol Treated Lactotrophs and Astrocytes, Measured with Amplex Red and Relative Normalized Changes in Astrocyte Lipid Fractions Measured with HPLC, Related to STAR Methods

| | Mean (% of control) | SEM (%) | p |
|---|---|---|---|
| **Lactotroph cholesterol – 10 mM MβCD for 30 minutes (Amplex Red) (n = 6)** | | | |
| CHOL | 56.79 | 10.62 | 0.008 |
| **Lactotroph cholesterol – 10 mM MβCD + cholesterol for 30 min (Amplex Red) (n = 6)** | | | |
| CHOL | 345.67 | 39.75 | <0.001 |
| **Astrocyte cholesterol – 10 mM MβCD for 30 minutes (Amplex Red) (n = 8)** | | | |
| CHOL | 52.76 | 4.85 | < 0.001 |
| **Astrocyte cholesterol – 10 mM MβCD + chol for 30 minutes (Amplex Red) (n = 8)** | | | |
| CHOL | 209.98 | 15.12 | < 0.001 |
| **Astrocyte lipid fractions – 10 mM MβCD for 30 minutes (HPTLC) (n = 3)** | | | |
| DOPE | 163.46 | 41.69 | 0.16 |
| DOPI | 109.42 | 20.49 | 0.65 |
| DOPS | 87.55 | 14.12 | 0.40 |
| DOPC | 102.62 | 10.02 | 0.8 |
| SM | 91.47 | 21.79 | 0.7 |
| CHOL | 59.60 | 11.85 | 0.027 |

### Table S3. Comparison of Unitary Exocytotic Vesicle Fusion in Lactotrophs and Astrocytes Recorded by the High-Resolution Membrane Capacitance Technique, Related to Figure 6.

| Measurement | Lactotrophs | | | Astrocytes | | |
|---|---|---|---|---|---|---|
| | REST | MβCD | MβCD − chol | REST | MβCD | MβCD + chol |
| Frequency. of reversible events (events/s), median (IQR) (no. of recordings) | 0.035 (0.023–0.052) (n = 52) | 0.007 (0.004–0.012)** (n = 66) | 0.001 (0.007–0.032)* (n = 55) | 0.029 (0.019–0.040) (n = 27) | 0.009 (0.006–0.014)** (n = 21) | 0.022 (0.015–0.027) (n = 27) |
| Frequency of irreversible events (/sec) | 0.003 (0–0.004) (n = 52) | 0.003 0.001–0.004) (n = 66) | 0.002 0–0.003) (n = 55) | 0.004 0.003–0.005) (n = 27) | 0.002 0.001–0.002)** (n = 21) | 0.003 0.002–0.005) (n = 27) |
| Open pore probability (%), median (IQR) (no. of recordings) | 1.79 (1.09–3.47) (n = 52) | 0.56 0.20–0.88)** (n = 66) | 0.86 0.34–1.87)* (n = 66) | 0.47 0.37–0.63) (n = 27) | 0.28 0.09–0.51)** (n = 21) | 0.58 0.34–0.81) (n = 27) |
| Fusion pore dwell time (s), median (IQR) (no. of recordings) | 0.30 (0.17–0.61) (n = 52) | 0.39 0.21–0.85)** (n = 66) | 0.30 0.16–0.74) (n = 55) | 0.11 0.06–0.27) (n = 27) | 0.18 0.07–0.61)** (n = 21) | 0.15 s 0.07–0.40) (n = 27) |
| % of reversible events with projection from Im to Re trace of admittance | 15% | 11% | 28% | 43% | 25% | 29% |
| Gp/Cv (pS/fF), median (IQR) (no. of events) | 10.0 (7.6–13.7) (n = 203) | 14.8 10.9–18.6)* (n = 44) | 4.8 4.4–5.0)** (n = 249) | 24.4 17.4–40.8) (n = 249) | 44.5 34.3–56.3)** (n = 31) | 27.2 16.7–50.7) (n = 133) |
| % reversible fusion (of all fusion events) | 94% | 74% | 90% | 91% | 84% | 87% |

The total recording times were 35.13 and 13.24 h in lactotrophs and astrocytes, respectively. *$p < 0.05$ versus REST; **$p < 0.01$ vs REST.

### A theoretical description of cholesterol-mediated fusion pore properties

To understand how cholesterol may affect the properties of the fusion pore, a water channel connecting the vesicle lumen (Figure 5A) with the extracellular space, we considered the relationship between the size of a secretory vesicle and the membranous fusion pore (Figure 5B), whereby the corresponding physical parameters are given. In this respect, relevant energy terms responsible for the invagination behavior have to be introduced.

The shape of a giant unilamellar phospholipid vesicle (GUV), whose size is comparable with that of an average biological cell, corresponds to the minimum of the membrane bending energy at a given area and volume (Deuling and Helfrich 1976; Seifert et al. 1991). Similarly, the energy of a small cell membrane invagination (i.e., secretory vesicle; Figure 5A) consists of the membrane bending energy and the attractive interaction energy between the secretory vesicle membrane and its lumenal environment. We assume that the contribution of the attractive interaction energy is proportional to the invagination surface area (i.e., secretory vesicle membrane area). The mechanical characteristics of the cell membrane invagination are determined by the membrane property of GUVs and, in addition, the energy contribution due to the different cholesterol areal densities between the membrane invagination (secretory vesicle membrane) and the plasmalemma enveloping the cell. Consequently, at the invagination boundary, i.e., at the fusion pore, line tension occurs.

Therefore, the mechanical energy of the invagination is given by the sum of three terms:

(1) the bending energy ($W_b$) is (Helfrich, 1973)

$$W_b = \frac{k_c}{2}\int (c_1 + c_2 - c_0)^2 dA \qquad \text{(eq. 1)}$$

where $c_1$ and $c_2$ are the membrane principal curvatures, $c_0$ is the membrane spontaneous curvature, $k_c$ is the membrane bending modulus, and $A$ is the area of the membrane invagination;

(2) the attractive interaction energy ($W_A$) at the invagination membrane takes the form

$$W_A = -wA \qquad \text{(eq. 2)}$$

where $w$ is the interaction energy per surface area;

(3) the energy of the invagination boundary ($W_L$) can be described by (Julicher and Lipowsky, 1996)

$$W_L = \eta L \qquad \text{(eq. 3)}$$

where $\eta$ is the line tension, which generates the tendency to constrict the fusion pore, and $L$ is the length of the circular boundary line.

Minimization of the invagination mechanical energy ($W_b + W_A + W_L$) leads to equations for the shape. We can neglect the effects concerning the pressure difference across the membrane ($\Delta p$) and lateral tension of the secretory vesicle membrane ($\lambda$). Namely, for the GUVs, typical dimensionless values of these parameters ($\Delta p R_{\mathrm{GUV}}^3/k_c$, $\lambda R_{\mathrm{GUV}}^2/k_c$, where $R_{\mathrm{GUV}}$ is the parameter equal to the radius of the sphere with the same area as the GUV $\sqrt{A_{\mathrm{GUV}}/4\pi}$), are in the order of magnitude equal to 1, which is comparable with the dimensionless value of the bending energy ($W_b/8\pi k_c$) (Svetina and Zeks, 1989). However, the sizes of secretory vesicles are about a hundred times smaller than GUVs. Therefore, neglecting the corresponding terms is justified. In addition, we assume that the reciprocal value of the spontaneous curvature ($1/c_0$) is the order of magnitude of the cell size, and thus $c_0$ can also be neglected. Consequently, the equations for the membrane midplane (Figure 5B) of the axisymmetric invagination can be written as (Julicher and Seifert, 1994)

$$\ddot{\psi}R = \frac{cos\,\psi\,sin\,\psi}{R} - \dot{\psi}cos\,\psi - \frac{\gamma}{2\pi k_c}sin\,\psi$$

$$\dot{\gamma} = \pi k_c\left(\frac{sin^2\,\psi}{R^2} - \dot{\psi}^2\right) + 2\pi w$$

$$\dot{R} = cos\,\psi \qquad \text{(eqs. 4)}$$

where $R$ (see Figure 5A) is the distance between the symmetry axis and the point on the membrane midplane of the invagination contour, $\psi$ is the membrane slope, $\gamma$ is the component of the transfer shear force in a radial direction, and the overdot denotes derivatives with respect to the arc length. The transfer shear force ($\gamma_{\text{pore}}$), proportional to the line tension ($\eta$) at the fusion pore,

$$\gamma_{pore} = 2\pi\eta \qquad \text{(eq. 5)}$$

is generating a radial force.

### Predictions of the theoretical model for the fusion pore shape

The procedure for obtaining the shape is as follows. For given values of the modulus ($k_c$), the interaction energy per surface area ($w$), and the initial membrane curvature at the invagination pole ($c_p$), the integration is performed by solving eqs. (4) from the invagination pole. The integration is halted at the invagination fusion pore, i.e., at the minimum of the radius $R$ (Figure 5A). The values for the parameters are tuned such that the radius of the fusion pore and the secretory vesicle membrane area assume their given values.

The results of the numerical integration are shown in Figures 5A and S6. For the sake of simplicity, we introduce the vesicle radius $R_0 = \sqrt{A/4\pi}$ (Figure 5A) to define the dimensionless quantities: the normalized radius of the membrane midplane at the fusion pore $r_{\text{Mpore}} = R_{\text{Mpore}}/R_0$, the normalized fusion pore radius $r_{\text{pore}} = R_{\text{pore}}/R_0$ (Figure 5B), the relative interaction $W = wR_0^2/2k_c$, and the radial force $\Gamma_{\text{pore}} = R_0\gamma_{\text{pore}}/8\pi k_c$, which contribute to fusion pore constriction. The difference between the radius of the membrane midplane at the fusion pore ($R_{\text{Mpore}}$) and the fusion pore radius ($R_{\text{pore}}$) corresponds to half of the membrane thickness ($h$), considered to be 2.5 nm (Kuchel and Ralston 1988), $R_{\text{Mpore}} = R_{\text{pore}} + h$.

The invaginations are essentially spherical (Figure 5A), and they can have different radii depending on the line tension. The results in Figure S6A show that the secretory vesicle shapes possess a maximum value of $W$ that is related to a change in $W_b$ with respect to the corresponding membrane area (Svetina and Zeks, 1989). At small values of $r_{\text{pore}}$, $W$ is small since at relatively small parts of the membrane, $c_1$ and $c_2$, which determine $W_b$, are large. In contrast, at the largest values of $r_{\text{pore}}$, $W$ is reduced because $c_1$ and $c_2$, and the relevant $W_b$, are small. Therefore, at these large fusion pore radii for a given value of $W$, two solutions exist (arrows, Figure S6A.) The prediction of how the dimensionless radial force depends on the normalized pore radius ( $r_{\text{Mpore}}$) is shown in Figure S6B. At small values of $W$, the radial force decreases with the relative interaction $W$, since reduction in the fusion pore radius results in an increase in $W_b$ (eq. 1).

### The role of the plasmalemmal cholesterol on the fusion pore shape

The effect of cholesterol areal density on the invagination behavior is related through the line tension. The cholesterol density appears higher in the secretory vesicle versus the plasma membrane (see Figure 1). We assume that the transfer shear force, determined by line tension (eq. 5) at the fusion pore, is proportional to the difference in cholesterol densities between the vesicle and the plasma membranes ($E_1 - E_0$). This feature can be conveniently described:

$$\gamma_{pore} = 8\pi k_c \alpha (E_1 - E_0) \quad \text{(eq. 6)}$$

where $\alpha$ is the model parameter. We also assume that the cholesterol density in the secretory vesicle membrane is larger at smaller fusion pore radii with respect to the secretory vesicle size. This can be approximately described by

$$E_1 = \frac{\beta R_0}{R_{pore}} \quad \text{(eq. 7)}$$

where $\beta$ is the model parameter. For chosen values of the parameters (β and $R_0$), the dependence of $E_1$ on $r_{M\text{pore}}$ is shown in Figure S6C. Inserting the expression for $E_1$ in eq. 6 leads to the dependence of the transfer shear force on the secretory vesicle at the fusion pore:

$$\gamma_{pore} = 8\pi k_c \alpha \left(\frac{\beta R_0}{R_{pore}} - E_0\right) \quad \text{(eq. 8)}$$

Considering eq. 8, $\gamma_{\text{pore}}$ is determined by $R_{\text{pore}}$. In Figure S6D, a dependence of the normalized pore radius $r_{\text{Mpore}}$ on the vesicle radius ($R_0$) is demonstrated for certain values of the parameters ($\alpha$, $\beta$, and $E_0$). At the smallest secretory vesicle radius (≲15 nm), the dependence may represent omega figures of fused synaptic-like vesicles seen on electron microscopy (Heuser, 1989), characterized by relatively wide fusion pores at small vesicle radii. At larger vesicle radii, the dependence represents relatively narrow fusion pores.

Figures S6E and 5D illustrate the dependence of the fusion pore radius ($R_{\text{pore}}$) as a function of the vesicle radius ($R_0$) for different values of the parameters, simulating the influence of the difference in cholesterol densities on line tension (the parameter $\alpha$), different plasmalemmal cholesterol levels (the parameter $E_0$), and the influence of the fusion pore radius (the parameter $\beta$). Figure S6E shows the corresponding dependencies for different values of the parameter $\alpha$ at given ratios between the parameters $\beta$ and $E_0$. These dependencies indicate that at large values of the parameter $\alpha$, its value is irrelevant. The corresponding interrelation between the fusion pore radius ($R_{\text{pore}}$) and vesicle radii ($R_0$) at large values of the parameter $\alpha$ can be obtained by setting the left side of eq. (8) to zero:

$$R_{pore} = \frac{\beta R_0}{E_0} \quad \text{eq. (9)}$$

The best fits to the experimental results are obtained considering eq. (9) (Figure S5). Figure 5D shows the corresponding dependencies for different ratios between the parameters $\beta$ and $E_0$ considering eq. (9). Note that the fusion pore radius ($R_{\text{pore}}$) for a given vesicle size ( $R_0$) at increased cholesterol density is relatively narrow (Figure 5D, high chol.), whereas at reduced cholesterol density, $R_{\text{pore}}$ is relatively wide (Figure 5D, low chol.).

**See the main text for references.**
